# Exon-Skipping Antisense Oligonucleotides for H3.3K27M-Altered Diffuse Midline Glioma Therapy

**DOI:** 10.64898/2026.04.03.715131

**Authors:** Lucia Yang, Qian Zhang, John E. Wilkinson, Adrian R. Krainer

## Abstract

Diffuse midline gliomas (DMGs) are a deadly class of pediatric high-grade brain cancers. Approximately 80% of pontine DMGs feature a dominant, somatic, heterozygous point mutation in the non-canonical histone H3.3-coding gene *H3-3A*. This dominant-negative mutation replaces lysine 27 with methionine (K27M) and prevents global K27 di- and tri-methylation of all wild-type histone H3 proteins. We aimed to target the H3.3K27M onco-histone pre-mRNA with splice-switching antisense oligonucleotides (ASOs) designed to promote skipping of *H3-3A* exon 2, as this constitutive exon comprises both the K27M mutation and the natural in-frame start codon of the gene. The lead ASO identified in a systematic screen specifically induced *H3-3A* exon 2 skipping, did not affect expression or splicing of the paralog gene *H3-3B*—which also encodes histone H3.3—and restored global H3K27me3 marks in patient-derived DMG cells grown as neurospheres. In a patient-derived orthotopic xenograft tumor mouse model, the lead ASO reduced proliferation and extended survival. Our results show the potential of exon-skipping ASOs targeting *H3-3A* exon 2 as a therapeutic option for H3.3K27M-altered DMG. More generally, they exemplify the strategy of using ASOs to induce skipping of a constitutive exon to effectively achieve gene downregulation.

## INTRODUCTION

Pediatric high-grade gliomas (pHGG) are a deadly class of pediatric brain cancers that account for 10-15% of all pediatric brain tumors^1–3^. pHGGs can be divided into four classes, each with its own unique localization, mutations, and outcomes^4^. One such class, driven by a lysine27-to-methionine (K27M) mutation in histone H3 (H3), is termed H3K27M-altered diffuse midline glioma (DMG) and occurs in midline structures, including the thalamus, pons, cerebellum, and spinal cord. H3K27M-altered DMG is primarily identified in younger pediatric patients, with peak incidence occurring in children 6 to 9 years old. These tumors are especially deadly, with a median survival after diagnosis of 10 months that can be attributed to the midline location and diffuse nature of these tumors, which limit clinical management by surgical resection and traditional chemotherapy. Radiation therapy is the mainstay palliative treatment option to provide symptom relief and delay disease progression^5^.

H3K27M mutations are observed in either of two genes encoding the non-canonical histone variants H3.1 and H3.3, respectively^6^. Mutations in the gene *H3C1* result in the mutant H3.1K27M histone, and are the rarer of the two K27M mutations. A heterozygous somatic A>T point mutation in the gene *H3-3A*, which encodes the non-canonical histone H3.3, is found in 70-80% of pontine DMGs^7–9^. This lysine-to-methionine substitution causes a global reduction in the levels of H3K27me3 in all histone H3 variants, including wild-type H3.1, H3.2, and H3.3 histones^10,11^. H3.3K27M binds and interferes with the EZH2 methyltransferase subunit of Polycomb repressive complex 2 (PRC2), explaining the global reduction of di- and tri-methylation on all histone H3 proteins^12^. This inhibition of PRC2-mediated H3K27me2/me3 marks also applies to the identical wild-type H3.3 protein expressed by the paralogous gene *H3-3B*. The emergence of H3K27M and consequent global loss of H3K27me3 are thought to be driving events in tumorigenesis. Two studies showed that knockdown via shRNA or knockout via CRISPR of H3.3K27M reduces tumor burden, extends latency, and restores differentiation programs^13,14^. In both studies, patient-derived glioma cell lines were treated in culture prior to establishment of patient-derived xenografts in mice.

We sought to use a potentially therapeutic method to downregulate H3.3K27M following xenograft establishment. Antisense oligonucleotides (ASOs) are synthetic nucleotide polymers that can be designed to bind any RNA target via Watson-Crick base-pairing and typically include chemical modifications that prevent nuclease degradation, improve cellular uptake and target binding, and block innate immune activation^15^. At present, 14 ASOs for conditions such as Duchenne muscular dystrophy and familial hypercholesterolemia have been approved by the U.S. Food and Drug Administration (FDA) or European Medicines Agency (EMA)^16–18^, and several others are in clinical trials for oncological disorders^19^. Importantly, two are approved for use in the central nervous system (CNS), with intrathecal delivery directly to cerebrospinal fluid (CSF) via lumbar puncture: nusinersen, a splice-switching ASO that induces *SMN2* exon 7 inclusion in spinal muscular atrophy (SMA) and tofersen, an RNase H-active gapmer ASO that degrades *SOD1* transcripts in an inherited form of amyotrophic lateral sclerosis (ALS).

We previously showed that the RNase H-active modality of ASO is effective at downregulating H3.3K27M oncohistone expression, extending survival in two murine models of H3.3K27M-altered DMG^20^. We aimed to expand this work with the pursuit of the other ASO modality, splice-switching. Although the genes coding for canonical histones are intronless, the *H3-3A* and *H3-3B* genes coding for the non-canonical histone H3.3 do contain introns, and thus their pre-mRNAs undergo splicing^21^. Here, we aimed to exploit this feature to develop a gene-specific splice-switching ASO targeting *H3-3A* to induce exon 2 skipping. Following ASO treatment, the exon-skipped mRNA lacks a start codon, preventing translation of the H3.3K27M histone. While this approach also targets wild-type *H3-3A*, leading to reduced wild-type H3.3, this is not expected to be detrimental, because *H3-3A*’s paralog, *H3-3B*, codes for the identical histone H3.3 and is ubiquitously expressed, and furthermore, the *H3f3a* and *H3f3b* orthologues in mice are genetically redundant^22,23^. Therefore, in the presence of wild-type *H3-3B*, knockdown of *H3-3A* is not anticipated to be harmful for non-tumor tissues. We thus aimed to develop a splice-switching ASO that induces *H3-3A* exon 2 skipping and downregulates mutant H3.3K27M, and to test its efficacy *in vitro* and *in vivo*.

## RESULTS

### Designing ASOs to Induce Skipping of *H3-3A* Exon 2

To induce *H3-3A* exon 2 skipping, we designed ASOs to target one of the essential elements for pre-mRNA splicing, the 5’ splice site. Sterically blocking this site should prevent binding of the U1 snRNP and subsequent spliceosome assembly and splicing of this constitutive exon^24^. Our goal was to target *H3-3A* in a gene-specific manner, sparing the paralog gene *H3-3B*. These two genes differ in overall size and exon-intron structure (**Figure S1A**), and although they both have an intron at the same position and in the same +1 phase of their first partially coding exon, there is substantial sequence divergence between them in and around these 5’ splice sites (**Figure S1B**). Thus, we performed a “micro-walk” with 17 ASOs positioned along the *H3-3A* exon 2 5’ splice site, with their target sites shifted in one-nucleotide “steps” (**Figure 1A**). The full list of ASOs (ASO58-74) tested is in **Supplemental Table 1**. All ASOs screened were 20-nucleotides long and modified with 2’-O-methoxyethyl (2’-MOE) groups, a phosphorothioate (PS) backbone, and methylated cytosines (5meC) (**Figure 1B**).

**Figure 1.**
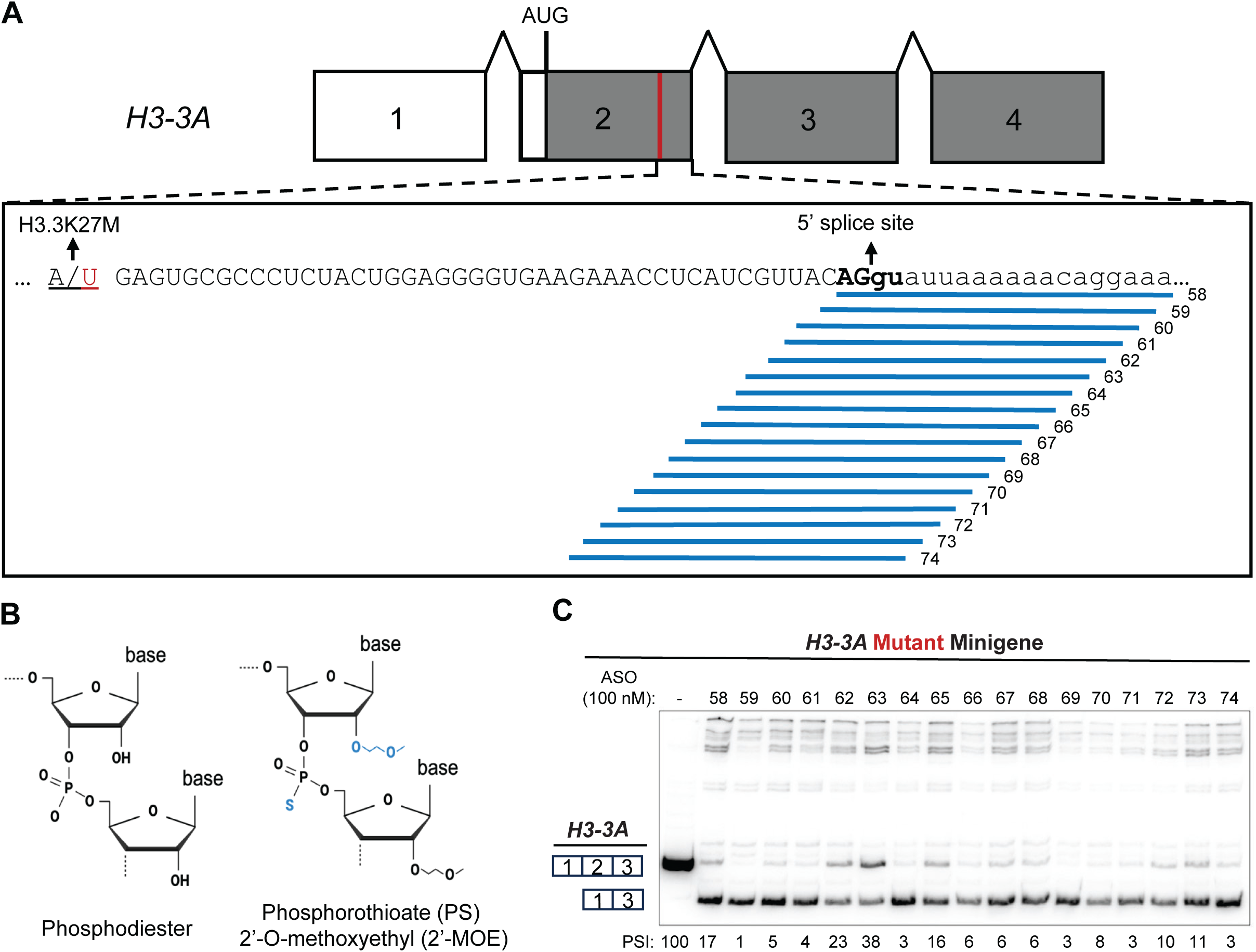
Micro-Walk Screen of ASOs Along the 5’ Splice Site of *H3-3A* Exon 2. (**A**) Seventeen ASOs were designed to target the 5’ splice-site of H3-3A exon 2. All ASOs were 20-mers, uniformly modified with 2’O-methoxyethyl (2’-MOE) groups, phosphorothioate (PS) linkages, and 5-methyl cytosines. The H3.3K27M mutation site is denoted in red. The 5’ splice site is bolded. Uppercase letters represent exonic sequences; lowercase letters represent intronic sequences. (**B**) Comparison of phosphodiester (PDE) bases with 2’-MOE-PS bases. Differences between PDE and 2’-MOE-PS bonds are denoted in blue. Created with BioRender.com. (**C**) ASO screen via co-transfection of 100 nM ASO and 200 ng mutant *H3-3A^K27M^* minigene plasmid in HeLa cells for 48 h. RT-PCR amplicons are derived from exogenous *H3-3A* mRNA through use of a forward primer specific to the minigene plasmid backbone. Percent spliced-in (PSI) with reference to *H3-3A* exon 2 is labeled below each sample.

We performed the ASO micro-walk by co-transfection with our previously described *H3-3A* minigene into HeLa cells^20^. The minigene comprises *H3-3A* exons 1, 2, and 3 with intact introns in a pcDNA3.1(+) plasmid vector. The vector also has a T7 promoter, such that we can amplify exogenous *H3-3A* by placing the forward primer in the T7 promoter and the reverse primer in exon 3, allowing us to capture RT-PCR amplicons with exon 2 included or skipped. All 17 ASOs screened induced *H3-3A* exon 2 skipping, compared to a non-treatment control of exogenous *H3-3A^K27M^*, with percent-spliced in (PSI) ranging from 1% (ASO59) to 38% (ASO63) at 100 nM ASO (**Figure 1C**). We confirmed the sequence of the skipped cDNA product by Sanger sequencing. Additional products with slower electrophoretic migration represent potential intron-retention and other mis-splicing events.

Next, we performed the micro-walk in a patient-derived glioma cell line, SU-DIPG-XIII, grown in neurosphere culture. This cell line was originally established from autopsy material from a patient diagnosed with H3.3K27M-altered DMG, formerly termed diffuse intrinsic pontine glioma^25^. We introduced the ASOs at 4 µM by free uptake (gymnosis), and again observed that all 17 ASOs induced *H3-3A* exon 2 skipping (**Figure S2**). We measured endogenous *H3-3A* mRNA with primers located in exon 1 and exon 3 to capture the full-length and exon-2-skipped isoforms. Based on the results of both micro-walk screens, we selected ASO59 (5’-TTCCTGTTTTTTAATACCTG-3’) as the lead splice-switching ASO.

### ASO59 Induced Dose-Dependent *H3-3A* Exon 2 Skipping in the Minigene Model

Following nomination of ASO59 as our lead splice-switching ASO, we performed a dose-response experiment to measure its potency in inducing *H3-3A* exon 2 skipping. We performed this with both wild-type *H3-3A^WT^* and mutant *H3-3A*^K27M^ minigenes, as ASO59 targets the 5’ splice site region, which does not overlap the mutation site. ASO59 induced *H3-3A* exon 2 skipping in both minigenes in a dose-dependent manner, with an EC_50_ of 9 nM for the wild-type minigene and 6 nM for the mutant *H3-3A*^K27M^ minigene (**Figure 2**). A scramble-sequence control ASO (ASO C1) of the same length and chemical modifications as ASO59 failed to induce exon 2 skipping, with the PSI remaining at 100%. We also tested an on-target, splicing-neutral control ASO (ASO C2) that binds to *H3-3A* exon 2 upstream of the H3.3K27M mutation, and it likewise failed to promote exon 2 skipping (**Figure S3A**). Furthermore, neither our control ASOs nor ASO59 induced detectable exon skipping or changed the levels of the *H3-3B* transcript. Sequences for ASOs C1 and C2 are listed in **Supplemental Table S1**.

**Figure 2.**
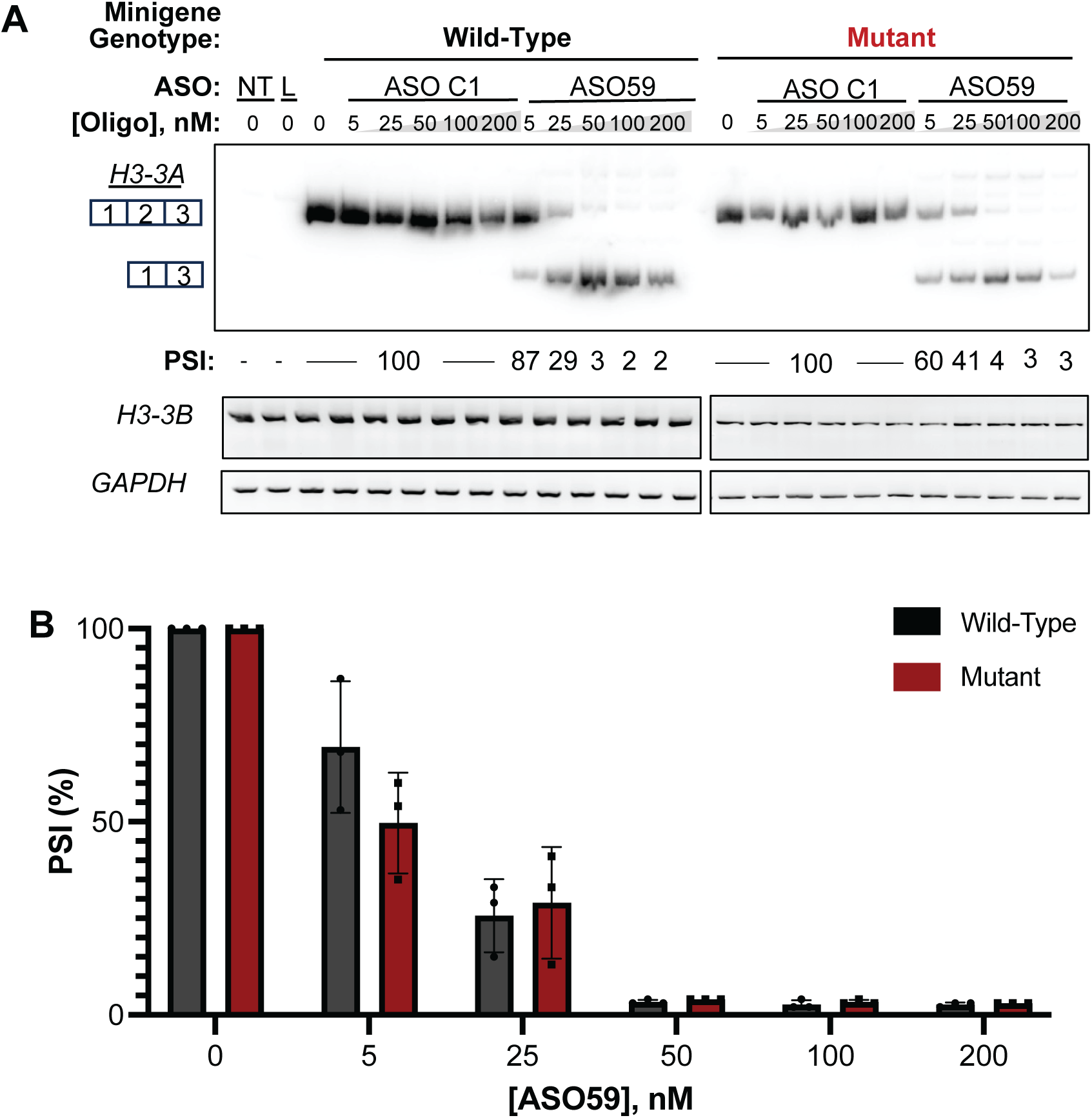
ASO59 Induces Dose-Dependent *H3-3A* Exon 2 Skipping in an *H3-3A* Minigene Reporter. (**A**) Representative RT-PCR gel of dose-response experiment with co-transfected wild-type *H3-3A* or mutant *H3-3A^K27M^* minigenes with scramble-sequence control ASO C1 or ASO59 in HeLa cells for 48 h. PSI is shown beneath each sample. (**B**) Quantification of PSI for each dose in each minigene (n=3). Each point represents one sample, with data represented as mean ± standard deviation. Non-treated control (NT), Lipofectamine-only control (L).

Our minigene was designed to express protein, as the *H3-3A* insert comprises the natural upstream Kozak sequence and start codon in *H3-3A* exon 2, plus an added in-frame stop codon at the end of *H3-3A* exon 3. The mutant minigene expressed H3.3K27M histone detectable by immunoblotting (**Figure S3B**). As expected, the protein expressed from the wild-type minigene did not react with the H3.3K27M-specific antibody. The product observed in the immunoblot is smaller than the endogenous 17-kDa histone H3.3, as the minigene lacks exon 4, which codes for the C-terminal portion of the protein. Co-transfection of the mutant *H3-3A* minigene and ASO59 led to reduced mutant H3.3K27M protein expression, relative to a mutant *H3-3A^K27M^* minigene-only control (**Figure S3C**).

### Validation of the Lead ASO in Additional Patient-Derived Cell Lines

Next, we tested ASO59’s effectiveness at inducing *H3-3A* exon 2 skipping in two additional patient-derived neurosphere lines kindly provided by Dr. Michelle Monje (Stanford University). In addition to SU-DIPG-XIII described above, we used a wild-type H3.3 glioma cell line, WT-H3.3, and another H3.3K27M-altered DMG cell line, SU-DIPG-XXIX^26^. We administered ASOs via free uptake at 1 µM and 4 µM doses for 48 h, and observed endogenous *H3-3A* exon 2 skipping in a dose-dependent manner with ASO59 administration, but not with ASO C1 administration (**Figure 3A**). Again, we verified that neither the ASO C1 nor ASO59 induced exon skipping in *H3-3B*. ASO59 induced *H3-3A* exon 2 skipping in all three cell lines, as expected, because ASO59 is gene-specific but not allele-specific.

**Figure 3.**
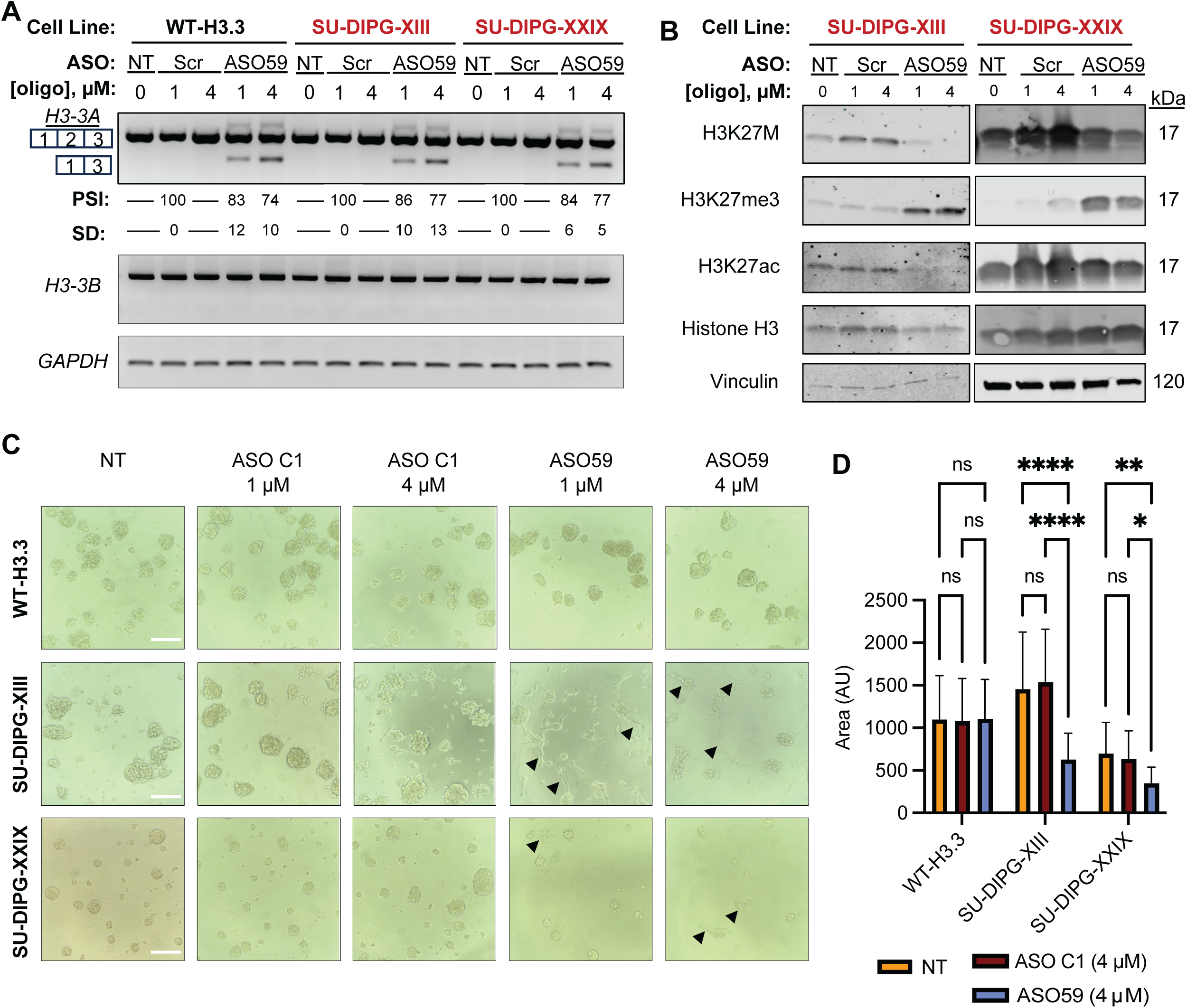
ASO59 Causes *H3-3A* Exon 2 Skipping, Restores H3K27me3, and Reduces Neurosphere Size in Patient-Derived Neurosphere Cell Lines. (**A**) RT-PCR gel of mRNA from the indicated patient-derived glioma cells grown as neurospheres and treated with ASO C1 (Scr) or ASO59 via free uptake for 5 d. Mean and standard deviation for each treatment are shown below (n=3). (**B**) Immunoblot of protein extracted from SU-DIPG-XIII and SU-DIPG-XXIX neurospheres following free uptake of ASO C1 (Scr) or ASO59 for 5 d. (**C**) Morphology of WT-H3.3, SU-DIPG-XIII, and SU-DIPG-XXIX patient-derived glioma cell lines following treatment with ASO C1 or ASO59 via free uptake for 5 d. Black arrowheads represent neurite-like outgrowth. Size bars represent 200 µm. (**D**) Quantification of average neurosphere area following no treatment (NT), free uptake of ASO C1 (4 µM), or free uptake of ASO59 (4 µM) for 5 d. Data represented as mean ± standard deviation. Neurosphere size differences between treatment groups were analyzed by ANOVA followed by pairwise comparisons using two-sample t-tests (\**p* < 0.05, \*\**p* <0.01, \*\*\**p* < 0.001, \*\*\*\**p* < 0.0001).

We next harvested protein lysates from WT-H3.3, SU-DIPG-XIII, and SU-DIPG-XXIX cells treated with the ASO C1 or ASO59 via free uptake. We confirmed that the WT-H3.3 cell line did not express H3.3K27M and did not exhibit any changes in global H3K27me3, H3K27ac, or total H3.3 expression following ASO treatment (**Figure S4**). In contrast, H3.3K27M was reduced and global H3K27me3 was restored following ASO59 treatment in mutant H3.3K27M-altered cell lines (**Figure 3B**). H3.3K27M expression was unchanged after ASO C1 treatment. In addition to measuring H3.3K27M and H3K27me3, we also stained for H3K27ac, as this histone modification was previously reported to be increased in mutant H3.3K27M-altered gliomas and is mutually exclusive with H3K27me3 marks^27^. ASO59 resulted in decreases in both H3.3K27M and global H3K27ac, with concomitant restoration of global H3K27me3 in ASO59-treated SU-DIPG-XIII and SU-DIPG-XXIX neurospheres (**Figure 3B**).

Following delivery of ASOs via free-uptake, the free-floating neurosphere morphology was unchanged in the wild-type H3.3 cell line; this result was expected, as the causative tumor-driving mechanism for this cell line is independent of H3.3K27M (**Figure 3C**). We also observed no difference in morphology or average neurosphere sizes in H3.3K27M-altered cell lines following no treatment or treatment with the ASO C1. In contrast, both SU-DIPG-XIII and SU-DIPG-XXIX exhibited neurite-like projections and significantly reduced neurosphere size following ASO59 treatment at 4 µM for five days (**Figure 3D**).

### ASO59 Inhibited Cell Proliferation and Cell-Cycle Progression

To characterize the cellular phenotypes following ASO59 administration in SU-DIPG-XIII, we performed a time-course proliferation assay using flow cytometry and observed that ASO59 at both 1 µM and 4 µM doses significantly reduced proliferation, compared to the non-treated and ASO C1-treated controls (**Figure 4A**). The proportion of viable cells remained similar between treatment groups at each timepoint, consistent with ASO59 resulting in reduced proliferation, as opposed to increased apoptosis (**Figure 4B**).

**Figure 4.**
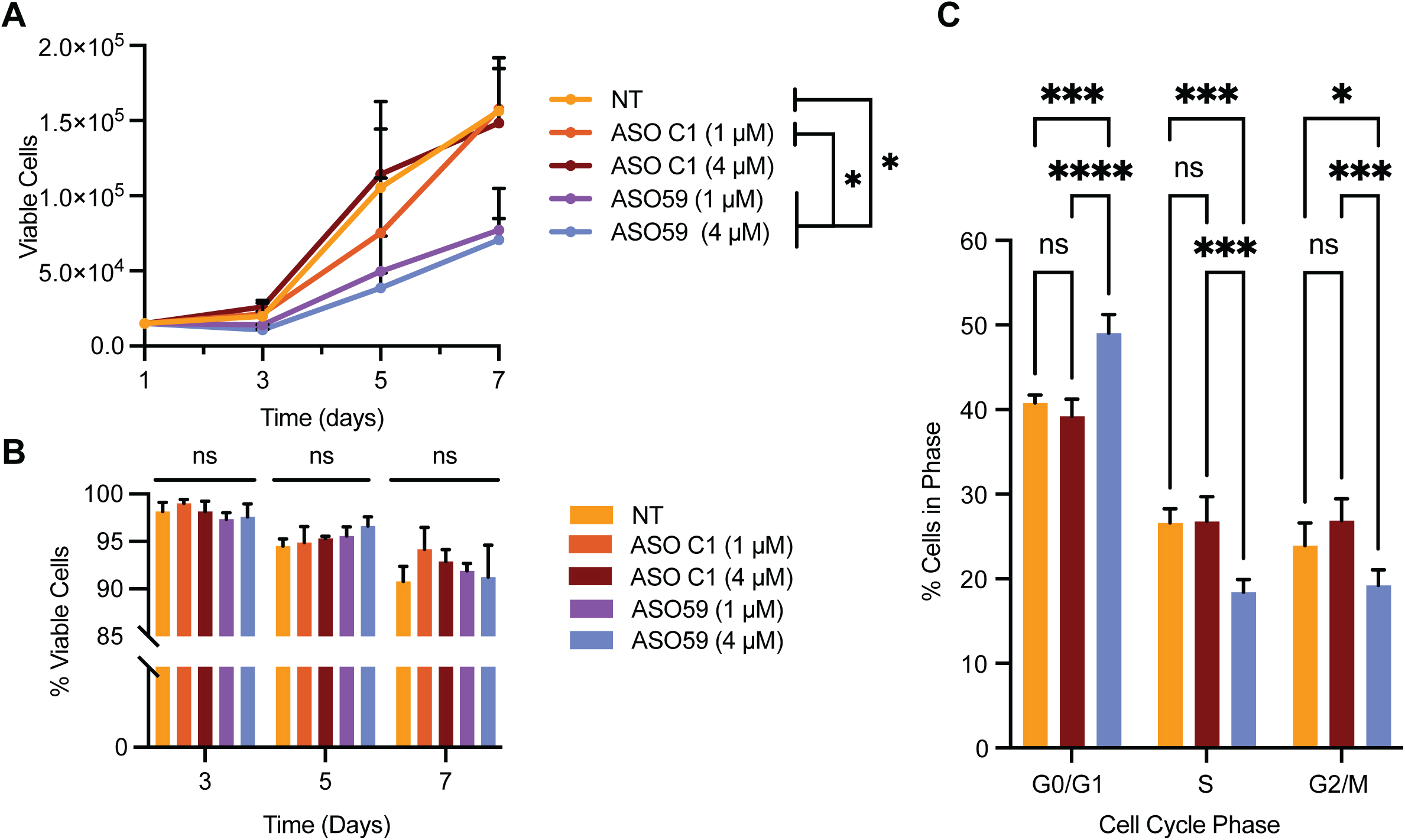
ASO59 Reduces Proliferation and Inhibits S-Phase Cell Cycle Entry in SU-DIPG-XIII. (**A**) Proliferation time-course of SU-DIPG-XIII cells following ASO C1 or ASO59 free uptake (n=3). (**B**) Percentage of viable cells at each time-point (n=3), analyzed by two-way ANOVA (factors: time, treatment). (**C**) Cell-cycle analysis of SU-DIPG-XIII cells following treatment with ASO C1 or ASO59 via free uptake on day 7 post-treatment (n = 3). Data represented as mean ± standard deviation. Proportion of cells in each cell-cycle phase analyzed by ANOVA followed by pairwise comparisons using two-sample t-tests (not significant (ns), \**p* < 0.05, ** *p* < 0.01, \*\**\*p* < 0.001, \*\*\*\**p* < 0.0001).

The decrease in SU-DIPG-XIII cell proliferation following ASO59 treatment, without a concomitant decrease in the percentage of viable cells related to the treatment, suggests inhibition of cell-cycle progression. It was previously reported that H3.3K27M inhibits the promoter of *P16/CDKN2A*, a tumor suppressor gene that slows cell division by acting as a checkpoint inhibitor at the G1-to-S phase of the cell cycle^28^. Thus, a larger proportion of H3.3K27M-altered cells are in S-phase, compared to WT-H3.3 cells. Here, we observed a similar effect in SU-DIPG-XIII cells, as untreated and ASO C1-treated cells exhibited a higher proportion of cells in S phase and G2 phase, compared to ASO59-treated cells (**Figure 4C**). Following ASO59 treatment, the proportion of cells in G0/G1 phase increased to a statistically significant degree, compared to non-treated and ASO C1-treated samples. Furthermore, the proportion of cells in S phase and G2/M phase decreased significantly following ASO59 treatment. We conclude that ASO59 treatment in SU-DIPG-XIII neurospheres limited proliferation through blocking cell-cycle progression, rather than by inducing extensive programmed cell death, as evident by the limited decrease in cell viability over time.

### Patient-Derived Xenograft (PDX) Model of H3.3K27M-Altered DMG

To study the effect of ASO59 in a preclinical mouse model of H3.3K27M-altered DMG, we established PDXs of SU-DIPG-XIII in NOD-*scid IL2rγ*^null^ (NSG) mice. The method for establishing PDXs from patient-derived glioma neurospheres was previously described^29^ and also employed in our lab^20^. We injected 50,000 luciferase-expressing SU-DIPG-XIII cells directly into the fourth ventricle/pons of postnatal day 1-2 mouse pups, and tracked tumor location and growth via bioluminescence imaging on postnatal day 21 (**Figure 5A**). We performed hematoxylin and eosin (H&E) staining and observed that the vehicle and ASO C1-treated tumors were highly proliferative and aggressive, with characteristic features including angiogenesis and pseudo-palisades surrounding areas of central necrosis (**Figure 5B**). We confirmed that our PDX of H3.3K27M-altered DMG recapitulated previously characterized features of DMGs, including expression of H3.3K27M that co-localizes with the human-specific marker hNCAM (**Figure 5C**), as well as high expression of the proliferative marker Ki67 and the oligodendrocyte precursor marker Olig2 (**Figure 5D**).

**Figure 5.**
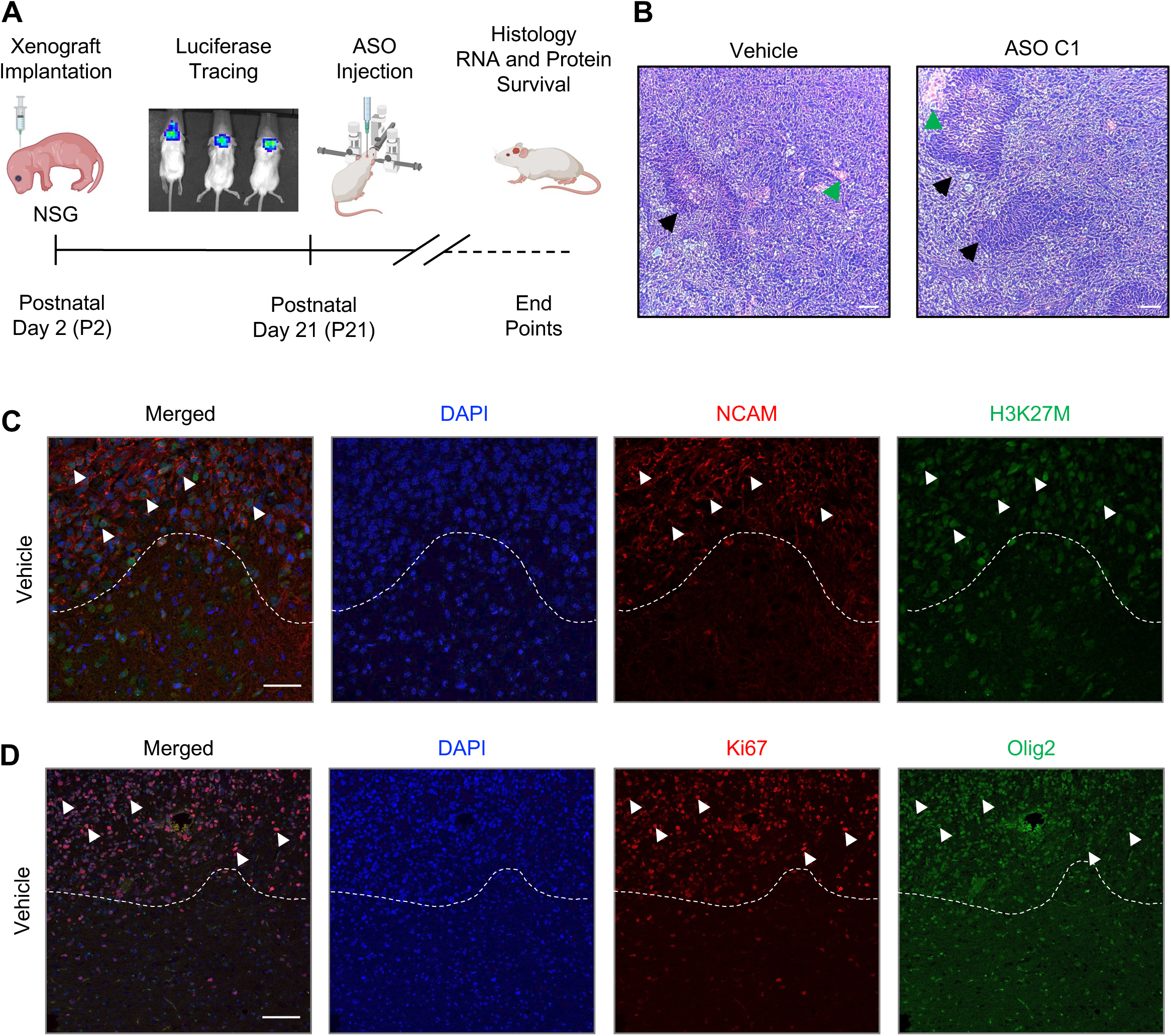
Patient-Derived Xenograft Model of H3.3K27M-Altered DMG. (**A**) Schematic of patient-derived xenograft tumor establishment using luciferized SU-DIPG-XIII cells in NSG mice. (**B**) Representative H&E stain of one vehicle-treated and one ASO C1-treated tumor, with black arrowheads indicating representative pseudo-palisades surrounding areas of central necrosis and green arrowheads indicating increased vascular proliferation. Images were taken at 10× magnification and the scale bar represents 50 µM. (**C**) Representative IF images of DAPI, human-specific marker hNCAM, and H3K27M from a vehicle-treated tumor. Representative co-labeling of NCAM and H3K27M indicated with white arrowheads. (**D**) Representative IF images of DAPI, proliferative marker Ki67, and glioma marker Olig2 in a vehicle-treated tumor. Representative co-labeling of Ki67 and Olig2 indicated with white arrowheads. IF images were taken at 20× magnification and scale bars represent 50 µm.

To visualize the ASO biodistribution in mouse brain tissue after delivery via intracerebroventricular (ICV) injection, we used an antibody, PS03, designed to detect the phosphorothioate backbone^30^. We validated this antibody in HeLa cells (**Figure S5A**) and SU-DIPG-XIII cells (**Figure S5B**), into which we delivered ASOs via transfection or free uptake for 24 hours. Following indirect immunofluorescence (IF) detection with co-staining for cytoplasmic tubulin and nuclear DAPI, we observed that both delivery methods resulted in all cells exhibiting cytoplasmic localization of the ASO, detectable with the PS03 antibody. We next performed IF staining in mouse brain tissues with established SU-DIPG-XIII xenografts, comparing samples that did or did not receive 200 µg ASO via ICV injection. We observed no signal with PS03 in the cortex and pons of the vehicle-treated sample, but observed a signal in both the cortex and pons of the ASO59-treated sample (**Figure S6**). Therefore, we conclude that ICV administration not only leads to ASO uptake in the target region of the pons and tumors, but also in the CNS as a whole in our PDX model.

### ICV Injection of ASO59 Induced *H3-3A* Exon 2 Skipping and Restored Global H3K27me3 in the PDX Mouse Model

Following validation of tumor establishment on postnatal day P21 by bioluminescence, we performed blinded 2-µL injections of either vehicle control (saline), ASO C1 (200 µg), or ASO59 (200 µg). After 7 days, we observed significantly reduced bioluminescence signals in the ASO59-treated tumors, relative to both the vehicle and ASO C1 controls (**Figure 6A**; p<0.001). Analysis of RNA extracted from the tumors revealed *H3-3A* exon 2 skipping in ASO59-treated tumors versus no skipping in ASO C1-treated tumors (**Figure 6B**). We also confirmed that *H3-3B* showed no differences in splicing or expression following control ASO or ASO59 treatment. Furthermore, we observed that H3.3K27M protein expression decreased and global H3K27me3 increased following ASO59 treatment (**Figure 6C**). Survival was significantly extended following ASO59 treatment, with the median survival extending from 26 days in vehicle- and ASO C1treated mice to 43 days in ASO59-treated mice (**Figure 6D**). Tumor samples were then analyzed in blinded fashion and assigned severity scores; severity scores ranged from 1 as least severe to 4 as most severe for infiltration, where less infiltration suggests slower growth and expansion. ASO59-treatment prevented tumors from reaching a severity score of 4, with a mean severity score of 2.0, compared to vehicle and ASO C1 treatment, which resulted in a higher proportion of tumors with greatest severity and a mean score of 2.57 (**Figure 6E**).

**Figure 6.**
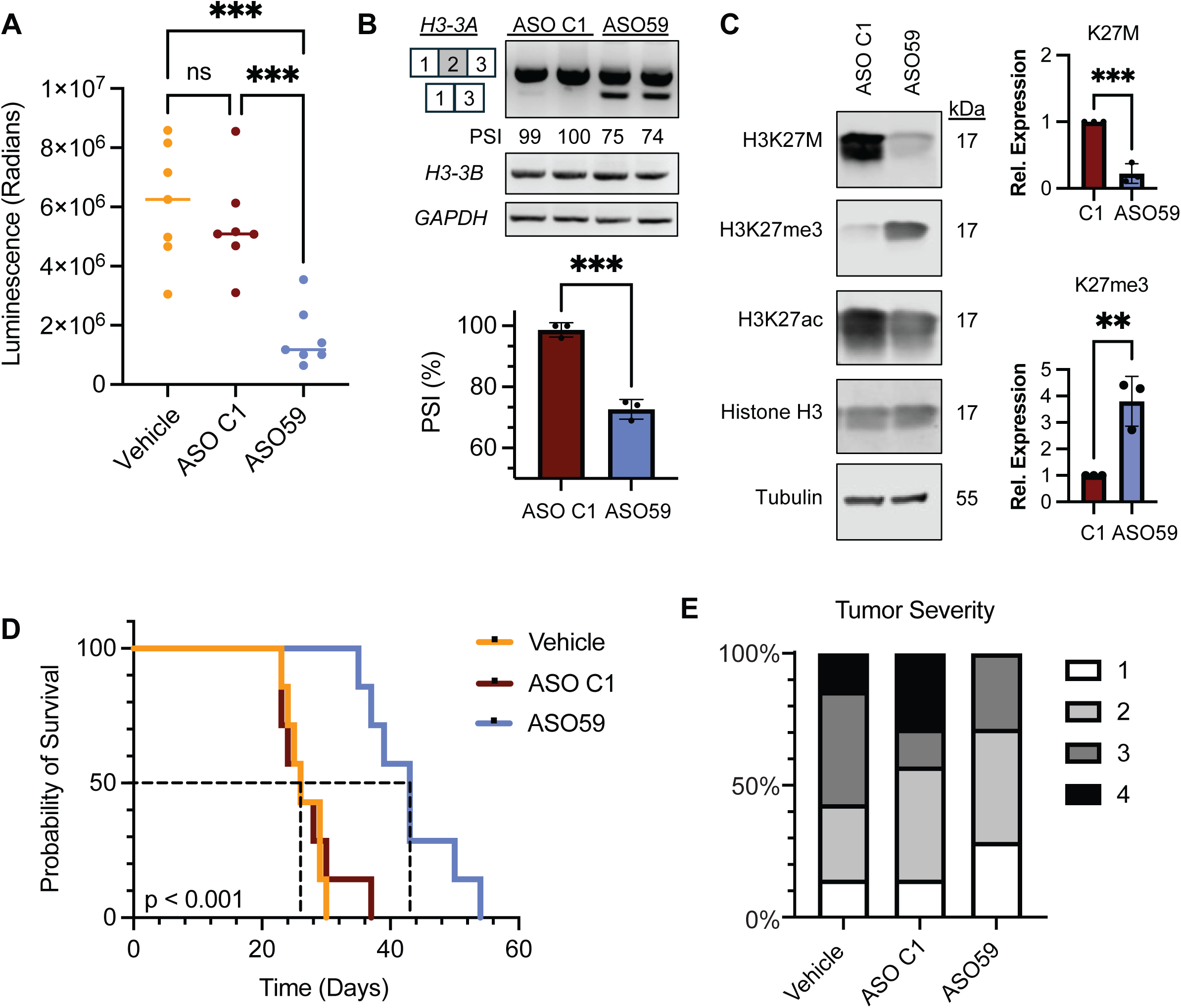
ASO59 Reduces Tumor Size, Induces *H3-3A* Exon 2 Skipping, Restores H3K27me3, and Extends Survival in a H3.3K27M-Altered DMG Patient-Derived Xenograft Model. (A) Comparison of bioluminescence (measured in radians) of vehicle-treated, ASO C1-treated, and ASO59-treated mice (n=7) at postnatal day 28. Each point represents one sample, with mean luminescence shown by a horizontal line. Measurements of bioluminescence were analyzed by ANOVA, followed by pairwise comparisons (\*\*\**p* < 0.001). (B) Representative RT-PCR gel and quantification of *H3-3A*, *H3-3B*, and *GAPDH* obtained from tumor samples treated with ASO C1 or ASO59. Percent spliced in (PSI) values are shown below each lane, and overall quantification of PSI mean ± standard deviation is shown in the bar graph. PSI was compared using Student’s t-test (n = 3, \*\*\**p* < 0.001). (C) Immunoblot of H3K27M, H3K27me3, H3K27ac, total histone H3, and loading-control tubulin from tumor tissues of ASO C1-treated or ASO59-treated mice. Relative expression mean ± standard deviation of H3K27M and H3K27me3 between ASO59-treated and ASO C1-treated samples (n=3) is graphed on the right and compared using Student’s t-test (\*\**p* < 0.01, \*\*\**p* < 0.001). (D) Kaplan-Meier curve of mouse cohorts following vehicle, ASO C1, or ASO59 ICV injection (n=7 for each treatment group). Median survival for vehicle-treated (26 d), ASO C1-treated (26 d), and ASO59-treated (43 d) groups is depicted with vertical dashed lines. Log-rank (Mantel-Cox) analysis was performed (*p* < 0.001). (E) Proportion of tumor-severity scores for tumors treated with vehicle control, ASO C1, and ASO59 (n=7 mice per treatment group), with 1 as least severe and 4 as most severe.

### ASO59 Reduced Proliferation and Induced Differentiation in the PDX Mouse Model

We next assessed the effect of ASO59 on proliferation and differentiation in our H3.3K27M-altered DMG PDX model. We performed IF on normal mouse tissue, ASO C1-treated tumors, and ASO59-treated tumors and we observed that Ki67 was absent in normal adjacent tissues, highly expressed in ASO C1-treated tumors, and downregulated in ASO59-treated tumors, relative to total cell number (**Figure S7**). We also performed H&E staining for all samples, again observing the classic histological features of high-grade gliomas, including high vascularization and pseudo-palisades surrounding areas of central necrosis in the vehicle control and ASO C1-treated samples (**Figure 7A**). We performed blinded histological-feature analysis for each tumor, which were based on the degree and extent of evidence of neural and glial cell differentiation, with a score of 1 being the least differentiated and a score of 4 the most differentiated. We observed that whereas all tumors were of histological grade IV, characteristic of most H3K27M-altered DMG tumors, the ASO59-treated tumors exhibited greater glial, neuronal, and overall differentiation compared to the vehicle-control and ASO C1-treated tumors (**Figure 7B**). Thus, from a histological perspective, ASO59 reduced proliferation and induced differentiation, as was evident from the downregulation of Ki67^+^-expressing cells and the histological grading, respectively.

**Figure 7.**
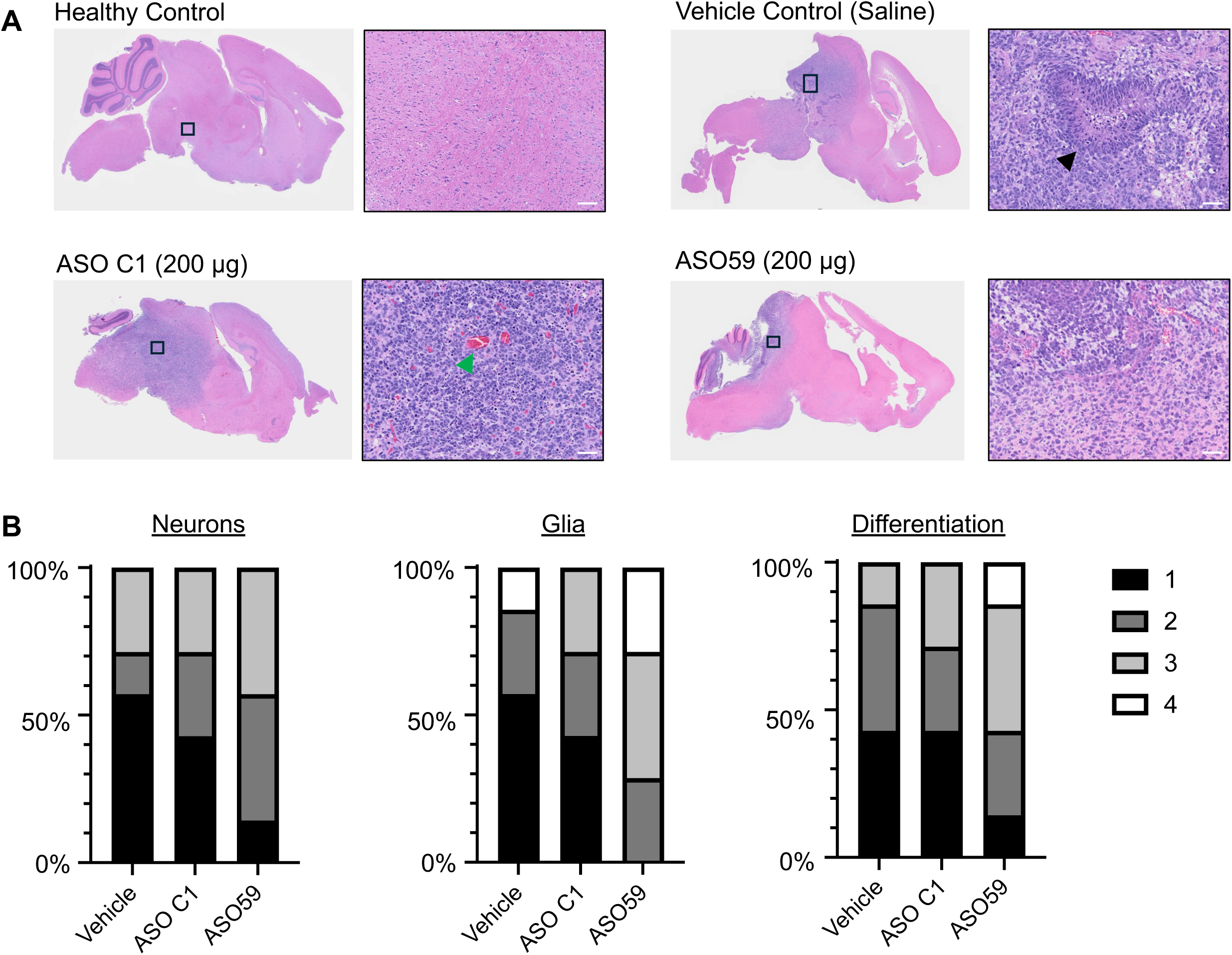
ASO59 Induces Differentiation of Glial and Neuronal Cells in a H3.3K27M-Altered DMG PDX Model. (**A**) Representative H&E staining of sagittal sections obtained from healthy control (top left), vehicle-treated (top right), ASO C1-treated (bottom left), and AOS59-treated (bottom right) mice. Representative regions within the pontine tumors are magnified to the right of each sagittal section, indicating pseudo-palisades surrounding central necrosis (black arrowhead) and vascular proliferation (green arrowhead). (**B**) Histological scoring of vehicle-treated, ASO C1-treated, and ASO59-treated tumors (n=7 mice per treatment group) for glial differentiation (left), neuronal differentiation (middle), and total differentiation (right) on a scale of 1 to 4, with 1 as absent to low and 4 as abundant.

## DISCUSSION

Here, we report the development of a splice-switching ASO designed to induce skipping of a constitutive exon and thus downregulate the oncohistone gene with the causative mutation in H3.3K27M-altered DMG. Our lead ASO, ASO59, targets the 5’ splice site of *H3-3A* exon 2, which contains both the eponymous H3.3K27M mutation and the natural start codon for the gene. We observed that ASO59-induced *H3-3A* exon 2 skipping consistently across our *H3-3A* minigene model, in multiple patient-derived DIPG cell lines, and in an orthotopic PDX mouse model.

The histone variant H3.3 is encoded by two paralogs, *H3-3A* and *H3-3B*. Although the proteins encoded by each gene are identical, the *H3-3A* and *H3-3B* nucleotide sequences are divergent, including at their respective exon 2 5’ splice sites, as well in their overall size and exon-intron structure (**Figure S1**). As a result, ASO59 specifically targets *H3-3A*, without inducing exon 2 skipping in *H3-3B*, thereby preserving wild-type histone H3.3 expression. Thus, the treatment should affect the tumor cells, which are dependent on H3.3K27M, but not normal tissues, which do not express the mutant oncohistone and continue to express the wild-type histone from the *H3-3B* paralog. Our approach underscores a broader therapeutic strategy of exon skipping to suppress translation of pathogenic alleles, especially in the case of dominant-negative mutations in genes that are either non-essential or are functionally compensated by paralogs.

Targeting the exon containing the major start codon of a gene for skipping can effectively reduce translation and is applicable in the context of H3K27M-altered DMG. The H3.3K27M mutation is a dominant-negative alteration that disrupts global K27 trimethylation across all the wild-type H3 histones. Prior studies demonstrated that knockdown or knockout of H3.3K27M globally restores the H3K27me3 marks^13,14^. In line with this, our lead ASO, ASO59, inhibited translation of the mutant H3.3K27M oncohistone and restored global H3K27me3 marks in patient-derived cell lines and a PDX model. Restoration of global H3K27me3 by ASO59 in turn reduced proliferation and prevented S-phase cell-cycle entry, shifting cells from S-phase and G2/M-phase to G1-phase. This result corroborates published work highlighting the role of mutant H3K27M as an inhibitor of an important cell-cycle checkpoint inhibitor, p16^28^. Finally, we observed reduced proliferation, increased differentiation, and prolonged survival following ASO59 treatment of a patient-derived xenograft model.

ASO59 significantly prolonged survival, though it did not eliminate tumor burden. This outcome mirrors previous findings with our lead “gapmer” ASO^20^, which likewise prolonged survival but did not achieve full tumor eradication. Several factors may contribute to this limitation, including failure to target all tumor cells, potential resistance mechanisms, or an ASO dosing regimen that is insufficient to overcome the aggressive growth kinetics or tumor burden of the particular H3K27M-altered DMG xenograft model. Nevertheless, our study demonstrates that a single dose of splice-switching ASO can effectively limit tumor growth and extend survival. Clinically, ASOs may offer a valuable resource in the toolkit to treat these aggressive tumors by helping reduce tumor growth to prevent a tumor-mass effect and keep treatment options such as radiation available. Or they may extend the therapeutic window in combination with other treatment modalities, such as GD2-targeting CAR-T cells currently in clinical trials^31^ or Modeyso (dordaviprone), a dopamine D2 receptor antagonist that was recently granted accelerated approval for treatment of adult and pediatric patients with recurrent H3K27M-altered DMG^32–34^. Furthermore, previous studies showed that radiosensitization of H3K27M-altered DMGs can be achieved through various mechanisms, including the use of DNA-damage repair inhibitors^35,36^ or modulation of mitochondrial metabolism^37^. It will be interesting to investigate the use of our lead ASOs, of either modality, in combination with focal radiation, to see if they can enhance radiosensitivity, limit tumor recurrence, or mitigate tumor-mass effects, such as hydrocephalus.

Our lab has now developed two lead ASOs—one splice-switching and one RNase H-active—to target *H3-3A* in H3.3K27M-altered DMG. Among the 13 clinically-approved ASOs currently in use, both modalities are well represented. Importantly, both modalities have been approved for use by intrathecal injection, exemplified by nusinersen for SMA and tofersen for *SOD1*-related ALS. The splice-switching ASO targeting *H3-3A* described here induces skipping of the exon harboring the start codon to prevent translation. This is a distinct mechanism from that of RNase H-active ASOs, which mediate transcript degradation through RNase H recruitment to the RNA-DNA duplex formed by the DNA-containing ASO and the target RNA. Because splice-switching ASOs are uniformly modified, they are expected to have longer half-lives than gapmers, and in theory, their risk of off-target effects is lower, because they do not directly trigger transcript degradation. Therefore, a splice-switching ASO with demonstrated efficacy for its intended splicing event may represent a safer therapeutic option.

We used intracerebroventricular injections to deliver ASOs to the in vivo PDX model. This route delivers the ASO directly into the CSF, allowing for ASO distribution to all CNS tissues via bulk flow^38^, followed by cell internalization, which we detected with a phosphorothioate backbone-specific antibody. ICV injection in mice, a stand-in for intrathecal injections in patients, provides a reliable solution to the inability of ASOs to cross the blood-brain barrier. Although some ASOs can result in sequence- and dose-dependent neurotoxicity^39^, such effects can be mitigated by formulation with divalent cations^40^.

We describe the use of one patient-derived glioma neurosphere line, SU-DIPG-XIII, to generate patient-derived xenografts. In the future, additional patient-derived xenografts should be tested to confirm the generality of our observations, and to compare ASO efficacy in different tumor contexts with different oncogene mutations, growth characteristics, and ASO-uptake properties. In this study, we employed a single 200-µg dose of ICV-administered ASO, which was the maximum tolerated dose for this chemistry in immunodeficient NSG adult mice. The CNS half-life of uniformly modified MOE ASOs is around 4-6 months^41,42^, which is longer than the mean 6-week survival we observed following ASO59 treatment in our PDX mouse model. However, in contrast to the terminally differentiated motor neurons targeted in SMA mice, some of the tumor cells in the DMG mouse model continue to proliferate, either because they failed to take up ASO, or because the internalized ASO is diluted out as the cells proliferate. Therefore, repeated ASO administration may enhance the anti-tumor effect, a possibility that should be tested in future studies, including dosing-schedule optimization.

In summary, we identified a lead splice-switching ASO that effectively induces *H3-3A* exon 2 skipping to reduce H3K27M mutant histone expression, restore global H3K27me3, and ameliorate tumorigenic phenotypes. This effect, observed in both patient-derived cell lines grown in cell culture and patient-derived xenografts in NSG mice, resulted in reduced proliferation and extended survival. We conclude that antisense-mediated *H3-3A* exon 2 skipping is sufficient to reduce translation of mutant H3.3K27M and rescue tumorigenic phenotypes induced by the mutant oncohistone (**Figure 8**). These findings support the continued development of splice-switching ASOs as a therapeutic strategy for H3K27M-altered DMGs, and more broadly for CNS disorders characterized by pathogenic gene expression. Our work also establishes an approach for gene downregulation via enforced skipping of a constitutive exon, expanding the therapeutic utility of splice-switching ASOs beyond their already-established applications in splicing correction, reading-frame restoration, and upregulation by redirecting splicing of non-productive mRNA isoforms^16, 43^.

**Figure 8.**
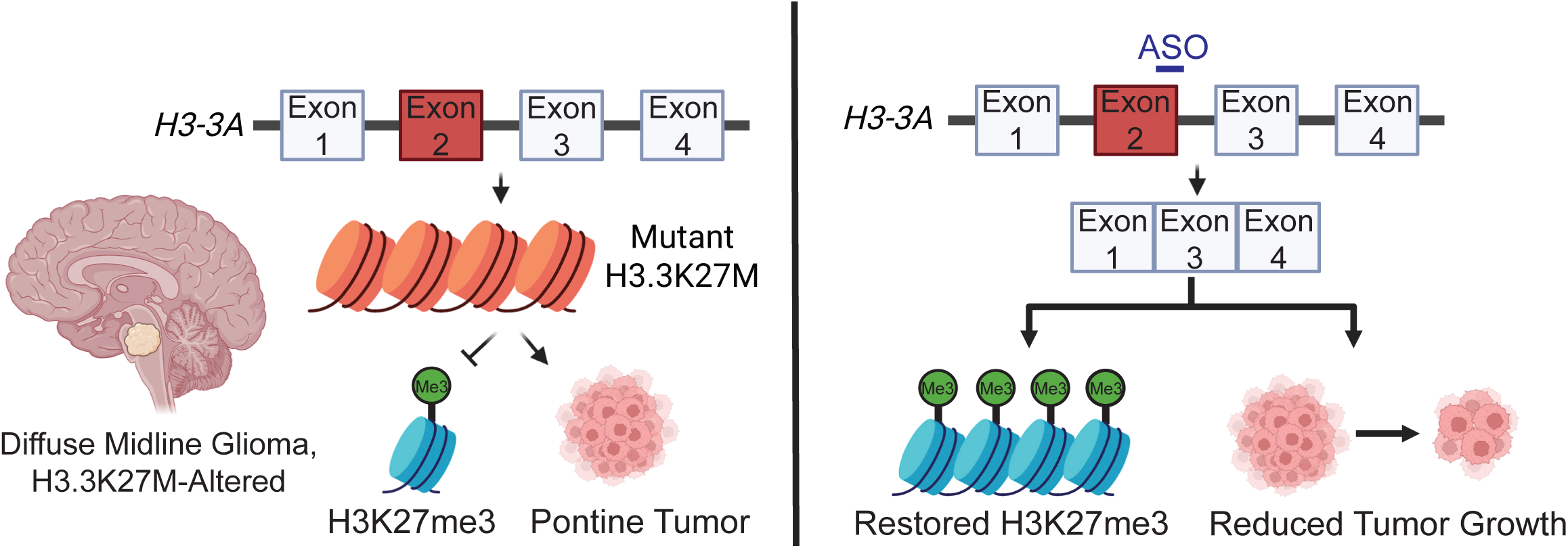
Model of Antisense-Targeted Skipping of *H3-3A* Exon 2 to Restore H3K27me3 and Reduce Proliferation in H3.3K27M-Altered Diffuse Midline Glioma Mouse Models. In H3.3K27M-altered diffuse midline gliomas (DMGs), mutant H3.3K27M, arising from an A>T point mutation in *H3-3A* exon 2, represses H3K27me3 expression and induces pontine tumor growth (left). A splice-switching ASO that induces *H3-3A* exon 2 skipping reduces H3.3K27M oncohistone translation, thereby restoring H3K27me3 marks and reducing tumor growth (right). Created with BioRender.com.

## DATA AVAILABILITY STATEMENT

All study data are included in the article and/or supplemental materials. Patient-derived neurosphere cell lines WT-H3.3, SU-DIPG-XIII, and SU-DIPG-XXIX were provided by Dr. M. Monje under an MTA with Stanford University.

## ACKNOWLEDGEMENTS

We thank Michelle Monje (Stanford University) for generously sharing patient-derived glioma cell lines, xenograft protocols, and advice. We thank Carl Ascoli (Rockland Immunochemicals) for generously sharing ModDetect reagents, protocols, and advice. This work was supported by grants from the Simons Foundation grant 552716, the V Foundation grant T2021-001, and NCI grant P01CA13106 Project 2 to A.R.K. L.Y. was supported by NIH NCI F30CA281196 and NIH NIGMS T32GM0084444 grants. We acknowledge assistance from the Cold Spring Harbor Laboratory Shared Resources, which are supported by NCI Cancer Center Support Grant 5P30CA045508.

## AUTHOR CONTRIBUTIONS

Q.Z. and A.R.K. conceived the project. L.Y., Q.Z., and A.R.K. designed the experiments. Q.Z. designed and performed the ASO micro-walk screen. L.Y. performed the *in vitro* validation and *in vivo* experiments. J.E.W. provided analysis of histological samples. L.Y. and A.R.K. prepared the manuscript draft. L.Y. and A.R.K. reviewed and edited the manuscript. A.R.K. provided resources and supervision. All authors have proofread and approved the manuscript for publication.

## SUPPLEMENTAL INFORMATION

Document S1. Figures S1-S7 and Tables S1, S2 and S3-S5. Worksheet S3.

Figure S1. Alignment of Human *H3-3A* and *H3-3B*. **(A)** Diagram of full transcripts of *H3-3A* and *H3-3B*. Rectangles represent exons and connecting lines represent introns. White boxes represent non-coding regions and black boxes represent coding regions. Base-pair length of exons is labeled above, and base-pair length of introns is labeled below. Total base-pair length and directionality are labeled below each transcript. **(B)** Sequence alignment of *H3-3A* and *H3-3B* exon 2 coding sequences. Nucleotides highlighted in grey represent mismatches. The red highlighted nucleotide denotes the site of the A>T point mutation of H3.3K27M. Nucleotides in uppercase represent exonic sequences; nucleotides in lowercase represent intronic sequences.

Figure S2. Microwalk of ASOs Targeting the 5’ Splice Site in a Patient-Derived Neurosphere Cell Line. RT-PCR gel of SU-DIPG-XIII patient-derived neurospheres treated with 4 µM of the indicated ASOs via free-uptake for five days. RT-PCR amplicons span endogenous *H3-3A* exon 1 to exon 3. PSI is labeled below each sample.

Figure S3. Validation of an Additional Control ASO and H3.3K27M-Specific Protein Expression in Minigenes. **(A)** RT-PCR of HeLa cells co-transfected with the *H3-3A* minigene and ASO C2, the on-target splicing-neutral control, for 48 h. PSI of each sample is labeled below. **(B)** Immunoblot of HeLa cells transfected with wild-type and mutant *H3-3A* minigenes for 48 h. (**C**) Immunoblot of HeLa cells transfected with 4 µg of wild-type *H3-3A*^WT^ or mutant *H3-3A*^K27M^ minigene and ASO59 for 48 h. Non-treated control (NT), Lipofectamine-only control (L).

Figure S4. ASO59 Treatment Does Not Affect a Wild-Type H3.3 Glioma Cell Line. Immunoblot of WT-H3.3, a wild-type H3.3 patient-derived neurosphere cell line, following free-uptake of a scramble-sequence control ASO C1 (Scr) or ASO59 for 5 d. Bands were normalized to vinculin and then normalized to non-treated controls. Fold change (FC) is labeled below each sample.

Figure S5. ModDetect PS03 Detects ASO59 Delivered via Transfection or Free Uptake in HeLa and SU-DIPG-XIII Neurosphere Cells. **(A)** IF of HeLa cells following transfection (50 nM) or free uptake (1 µM) of ASO59 for 24 h. Arrowheads highlight the ModDetect PS03 signal. (**B**) IF of SU-DIPG-XIII neurosphere cells following a 24-h incubation with ASO59 delivered by transfection (50 nM) or free uptake (1 µM). Images are at 20× magnification with zoomed-in frames split into each individual channel.

Figure S6. Phosphorothioate Groups in ASO59 Are Detectable Epitopes by ModDetect in Mouse Brain Tissues. IF of mouse brain tissues following ICV injection with saline (top half) or 200 µg of ASO59 (bottom half). Tissue sections from the cortex and pons from each sample are depicted. Images were taken at 20× magnification and the scale bar represents 50 µm.

**Figure S7. ASO59 Reduces Ki67 and H3K27M Expression in a Patient-Derived Xenograft Model of H3.3K27M-Altered DMG. (A)** Representative IF staining of DAPI, Ki67, and H3K27M in tissue sections from normal, ASO C1-treated, and ASO59-treated mice. The dashed line indicates the border between tumor and normal adjacent regions. Images are at 20× magnification and scale bars represent 50 µm. (**B**) Quantification of the ratio of Ki67+ cells out of total DAPI+ cells in tumors from ASO C1-treated and ASO59-treated mice. Data are represented as mean ± SD (n = 3; Student’s t-test, \*\*\**p* < 0.001).

Table S1. List of ASOs. ASO58 to ASO 74 were designed to target the 5’ splice site of *H3-3A* exon 2 and used for the microwalk experiments. Control ASOs include a scramble-sequence control for ASO59 (ASO C1) and an on-target splicing-neutral control that binds to *H3-3A* exon 2 upstream of the start codon (ASO C2).

**Table S2. List of Primers.**

Table S3. *In Vivo* Data from H3.3K27M-Altered DMG PDX Samples.

**Table S4. Summary Statistics of H3.3K27M-Altered PDX Survival.**

**Table S5. Survival Analysis of H3.3K27M-Altered PDX.**

## MATERIALS AND METHODS

### Antisense Oligonucleotides

Antisense oligonucleotides (ASOs) were purchased from Integrated DNA Technologies. For use in transfection or free-uptake experiments in cell culture, ASOs were resuspended in nuclease-free water to a concentration of 100 µM. For *in vivo* mouse injections, ASOs were resuspended in sterile saline (VetOne). ASO concentration was confirmed by spectrophotometry and the ASO stock was stored at −20 °C for short-term use and at −80 °C for long-term storage.

### Minigene

The wild-type *H3-3A^WT^* and mutant *H3-3A^K27M^*minigenes used here were previously described^20^. In brief, they comprise *H3-3A* exons 1, 2, and 3 with intact introns in a pcDNA3.1(+) vector. Minigene plasmids were amplified by transformation of One Shot^TM^ Stbl3^TM^ Chemically Competent *E. coli* (Thermo Fisher Scientific) and cultured in Luria Broth (LB) with ampicillin. Plasmids were purified with QIAGEN Plasmid Maxi Kits, resuspended in nuclease-free water, and stored at −20 ^°^C for short-term use and at −80 ^°^C for long-term storage.

### Cell Culture

HeLa cells were obtained from the Cold Spring Harbor Laboratory Cell Culture Core. Cells were cultured in Dulbecco’s-Modified Eagle Medium (DMEM, Gibco) supplemented with 10% fetal bovine serum (FBS) and 1% penicillin-streptomycin, and grown in 5% CO_2_ at 37 °C. Cells were passaged when they reached 90% confluency by treatment with 1× trypsin.

### Culture of Patient-Derived Cells

De-identified patient-derived cells were a generous gift from Dr. Michelle Monje (Stanford University). Cell lines were cultured as neurospheres as previously described^25^. In brief, tumor stem medium (TSM) base containing Neurobasal-A (Gibco), Dulbecco’s Modified Eagle Medium Nutrient Mixture F12 (DMEM/F12, Gibco), penicillin-streptomycin, GlutaMAX, HEPES buffer, sodium pyruvate, and MEM non-essential amino acids was prepared and filtered through a 0.2 µm filter. Prior to use, TSM was supplemented with human PDGF-AB (20 ng/mL; PeproTech), human EGF (20 ng/mL; PeproTech), human FGF (20 ng/mL; PeproTech), and 50× B-27 Supplement Minus Vitamin A (Thermo Fisher Scientific; 1:50) growth factors. Cell lines were luciferized by transduction (pLenti PGK V5-LUC Neo (w623-2); Addgene, plasmid #21471) and selected with G418 (gentamicin, 1 µg/mL) for one week. All cells were incubated in 5% CO_2_ at 37 °C. All cell lines were frozen and stored in BAMBANKER freezing medium (Fujifilm Biosciences) at −80 °C.

### ASO Transfection and Free Uptake

For HeLa cell transfection, cells were seeded at 50% confluency. After 24 h, ASO and/or plasmid constructs were transfected with Lipofectamine 2000 (Thermo Fisher Scientific) per the manufacturer’s recommended protocol. Cells were harvested for RNA and protein after 48 h. For patient cells, tissue-culture plates and wells were prepared with Matrigel (Corning) and seeded for 24 h. ASOs and/or plasmid constructs were transfected with either Lipofectamine 3000 (Thermo Fisher Scientific) or Lipofectamine RNAiMAX (Thermo Fisher Scientific) for 48 to 72 h, using the manufacturer’s recommended protocol. For free-uptake experiments, cells were seeded as free-floating neurosphere cultures, and ASOs were added after 1 h. ASOs and media were replenished on day 3 and cells were harvested on day 5.

### RNA Extraction and RT-PCR

Cell culture and tissue samples were harvested with TRIzol Reagent (Thermo Fisher Scientific) as per the manufacturer’s recommendation. RNA samples were reverse-transcribed using ImProm-II™ Reverse Transcriptase (Promega) as per the manufacturer’s recommended protocol. RNA and cDNA were stored at −20 °C for short-term use and at −80 °C for long-term storage. cDNA samples were prepared for PCR using Q5 High-Fidelity Polymerase (New England Biolabs) or Ampli-Taq Polymerase (Thermo Fisher Scientific). Radioactive PCRs were conducted with the incorporation of radioactive ɑ-^32^P-dCTP (Perkin-Elmer). All primers used are listed in **Supplemental Table S2**.

### Immunoblotting

Cell culture and tissue samples were harvested in RIPA Lysis Buffer (10 mM Tris pH 8.0; 140 mM NaCl; 1% Triton X-100; 0.1% sodium deoxycholate; 0.1% sodium dodecyl sulfate; 1 mM EDTA, 0.5 mM EGTA). Protein concentrations were determined using Bradford reagent. Protein samples were mixed with 4× SDS Loading Dye (20% 1M Tris HCl, pH 6.8; 0.08% sodium dodecyl sulfate, 40% glycerol, 4% beta-mercaptoethanol, 0.01% bromophenol blue), run on 8-16% SDS-polyacrylamide gels, transferred onto nitrocellulose membranes, and then blocked with 5% BSA in 0.01% Tween-20 in TBS. Primary antibodies were added and incubated overnight at 4 °C. Secondary antibodies were incubated for 1 h at room temperature and included the following:: IRDye® 680RD Goat Anti-Rabbit IgG Secondary Antibody (1:10,000; LI-COR, 926-68071) and IRDye® 800RD Goat Anti-Mouse IgG Secondary Antibody (1:10,000; LI-COR, 926-68070). Membranes were imaged with a LI-COR Odyssey and images were analyzed with ImageJ. Membranes were stripped with Stripping Buffer (LI-COR) as needed. Primary antibodies included anti-H3K27M (1:1000; rabbit; Millipore Sigma, ABE419), anti-H3K27me3 (1:1000; rabbit; Cell Signaling, C36B11), anti-H3K27ac (1:1000; rabbit; Abcam, ab4729), anti-histone H3 (1:1000; rabbit; Abcam, ab1791), and anti-vinculin antibody (1:1,000; mouse; Santa Cruz Biotechnology, sc-73614).

### Neurosphere Microscopy

Brightfield images of neurospheres were acquired with a Revolve Microscope (ECHO). All images were at 10× magnification, and size bars represent 200 µm. Neurosphere area was measured using ImageJ.

### Proliferation Assay

Cells were cultured in growth factor-free TSM medium for 24 h. Cells were then seeded as free-floating cell cultures and ASOs were added to the medium after 1 h. Media and ASOs were replenished on day 3. For each day of proliferation measurements, cells were resuspended into single-cell suspension with Accutase (Thermo Fisher Scientific), and Guava® ViaCountTM (Cytek Biosciences) was added. Cell viability and proliferation were measured with a Cytek® Guava® easyCyte™ Flow Cytometer. Data were analyzed with FlowJo.

### Cell Cycle

Cells were cultured in growth-factor-free TSM medium for 24 h. Cells were then seeded as free-floating cell cultures and ASOs were added 1 h post-seeding. Media and ASOs were replenished on day 3. On day 5, cells were resuspended in a single-cell suspension. Equal amounts of each treatment group were then incubated in 70% ethanol overnight. Cells were then resuspended in Guava® Cell Cycle Reagent (Cytek Biosciences) for 1 h. Cell-cycle analysis was performed with a Cytek® Guava® easyCyte™ Flow Cytometer. Data were analyzed with FlowJo.

### Generation of Patient-Derived Xenograft (PDX) Mouse Model of H3K27M-Altered Diffuse Midline Glioma

NOD-SCID-Gamma (NSG) mice were obtained from The Jackson Laboratory (strain #005557). Mice were housed according to CSHL IACUC guidelines. Newborn pups at postnatal day 1-2 were injected with 50,000 patient-derived cells using a Hamilton syringe and 30-gauge needle directly into the pons (3 mm posterior to the lambda suture, 3 mm deep). At postnatal day 21, pups were weaned, and tumor development was confirmed with a Xenogen IVIS Imager. Mice were injected with D-luciferin (Gold Bio, Catalog #LUCK-100) intraperitoneally, as per manufacturer’s recommendation. Mice with similar-size tumors were then randomly sorted into treatment groups, maintaining the same number of male and female mice.

### ASO Injection in Mice

Surgeries and post-surgical care were performed according to CSHL IACUC guidelines. Treatment administration was blind with respect to lead ASO and control ASO. Mice were anesthetized with a ketamine and dexmedetomidine cocktail. Prophylactic buprenorphine analgesic was injected subcutaneously. Mice were loaded into the stereotaxic apparatus and a maximum of 3 µL of ASO was injected as a single dose into the fourth ventricle (coordinates: 1 mm behind lambda suture, 2 mm deep). Mice were then administered Antisedan/atipamezole and allowed to recover fully. Mice were checked daily following surgery for post-surgical analgesia and wound care.

### Immunofluorescence in Cell Culture

Cells were processed for immunofluorescence staining 24 h after transfection or gymnosis. All washes were performed with 1× phosphate buffered saline (PBS). Cells were fixed with 4% paraformaldehyde, then permeabilized with 0.1% Triton X-100 in PBS. Cells were then blocked in 1% bovine serum albumin (BSA) in PBS. Primary antibodies were incubated overnight at 4 °C and included anti-alpha-tubulin (1:200; rabbit; Cell Signaling, 2125) and ModDetect PS03 (1:1000; mouse; Rockland Immunochemicals). Secondary antibodies were incubated for 1 h at room temperature and included the following: Goat anti-Mouse IgG (H+L) Cross-Adsorbed Secondary Antibody, Alexa Fluor^TM^ 555 (1:10,000; Thermo Fisher Scientific, A32727) and Goat anti-Rabbit IgG (H+L) Cross-Adsorbed Secondary Antibody, Alexa Fluor^TM^ 488 (1:10,000; Thermo Scientific, A32731). Cells were cross-stained with DAPI and then mounted onto slides with ProLong Gold^TM^ Antifade Mountant (Invitrogen^TM^).

### Immunohistochemistry and Immunofluorescence of Mouse Tissues

Tissues were dissected from mice post-euthanasia and fixed in 4% para-formaldehyde overnight. Brains were then transferred to 1× PBS and sliced into sagittal sections and paraffin-embedded. Tissue sections were deparaffinized with successive xylene and alcohol washes. H&E staining was performed by the CSHL Histology Core and stained sections scanned with an Olympus VS200 microscope. All H&E sections were evaluated by a histopathologist (J.E.W.) blinded to treatment, and scored for features from a scale of 1 (lowest) to 4 (highest), relative to other tumors in the set (**Supplemental Table S3**).

For immunofluorescence, antigen retrieval was performed with a heat-induced citrate buffer method. Tissue sections were blocked with Background Buster (Innovex Biosciences). Primary antibody was incubated overnight at 4 °C. Primary antibodies included anti-H3K27M (1:1000; rabbit; Millipore Sigma, ABE419), anti-Ki67 (1:50; mouse; BD, 550609), anti-NCAM (1:200; mouse; Santa Cruz Biotechnology, sc-106), and anti-Olig2 (1:500; rabbit; Millipore, AB9610). Secondary antibodies (described above) were incubated for 1 h at room temperature. Following washes, the slides were counterstained with DAPI and mounted with coverslips. Confocal microscopy images were taken with a Zeiss LSM 780 confocal microscope at 10×, 40×, and 63× magnification. Images were processed and analyzed with ZEISS ZEN software and ImageJ.

### Statistical Analysis

All data are presented as the mean ± standard deviation. All statistical analyses were performed with GraphPad Prism 10. Comparisons of continuous variables between treatment groups were analyzed using Student’s t-test. Cell viability was analyzed with 2-way ANOVA with Geisser-Greenhouse correction and Tukey’s post-hoc tests. Neurosphere area was analyzed with ANOVA followed by pairwise comparisons using Student’s t-test. Mouse survival data were analyzed using the Kaplan-Meier method and the log-rank (Mantel-Cox) and Gehan-Breslow-Wilcoxon tests (**Supplemental Table S4 and S5**). P-values < 0.05 were considered statistically significant with values denoted as follows in the figures: \**p* < 0.05, \*\**p* < 0.01, \*\*\**p* < 0.001, \*\*\*\**p* < 0.0001.

