## Supplementary material for "Exon-Skipping Antisense Oligonucleotides for H3.3K27M-Altered Diffuse Midline Glioma Therapy": Document S1

Figure S1

**A**

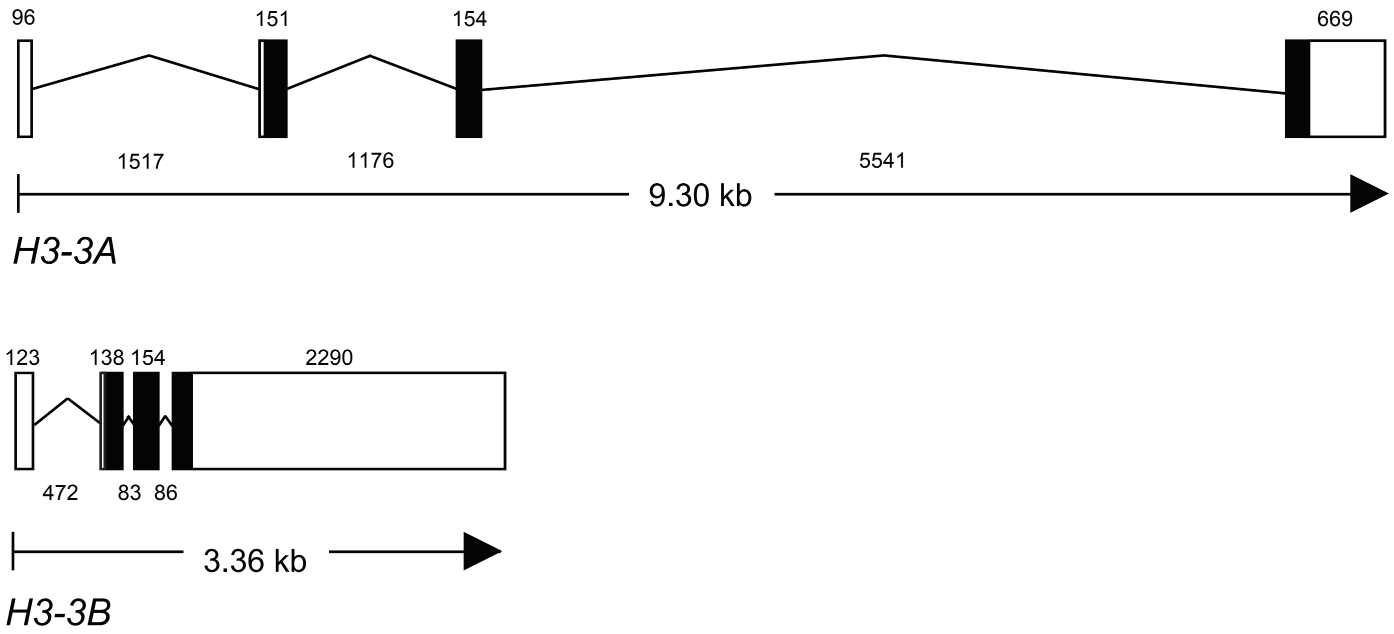


**B**

```
Exon 2 Alignment
H3-3A    ...ATGGCTCGTACAAAGCAGACTGCCCGCAAATCGACCGGTGGTAAAGCAC
H3-3B    ...ATGGCCCGAACCAAGCAGACTGCTCGTAAGTCCACCGGTGGGAAAGCCC
Protein  ...-M--A--R--T--K--Q--T--A--R--K--S--T--G--G--K--A--

H3-3A    CCAGGAAGCAACTGGCTACAAAAGCCGCTCGCAAGAGTGCGCCCTCTACTGG
H3-3B    CCCGCAAACAGCTGGCCACGAAAGCCGCCAGGAAAAGCGCTCCCTCTACCGG
Protein  P--R--K--Q--L--A--T--K--A--A--R--K--S--A--P--S--T--G

H3-3A    AGGGGTGAAGAAACCTCATCGTTACAGgtattaaaaaacaggaaaaaaatgg
H3-3B    CGGGGTGAAGAAGCCTCATCGCTACAGgtaggtcggcggggaacaatggc
Protein  --G--V--K--K--P--H--R--Y--R
```

Figure S2

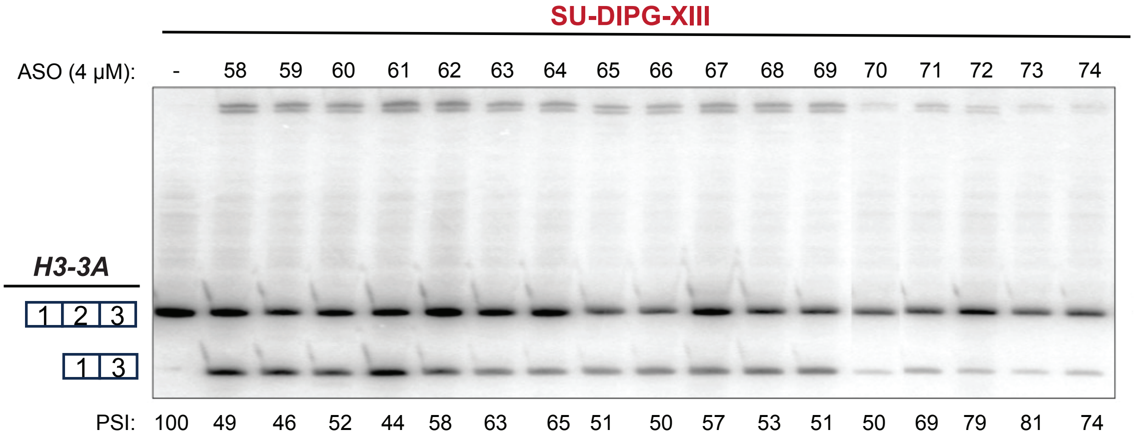

SU-DIPG-XIII
ASO (4 µM): - 58 59 60 61 62 63 64 65 66 67 68 69 70 71 72 73 74
H3-3A
1 2 3
1 3
PSI: 100 49 46 52 44 58 63 65 51 50 57 53 51 50 69 79 81 74

Figure S3

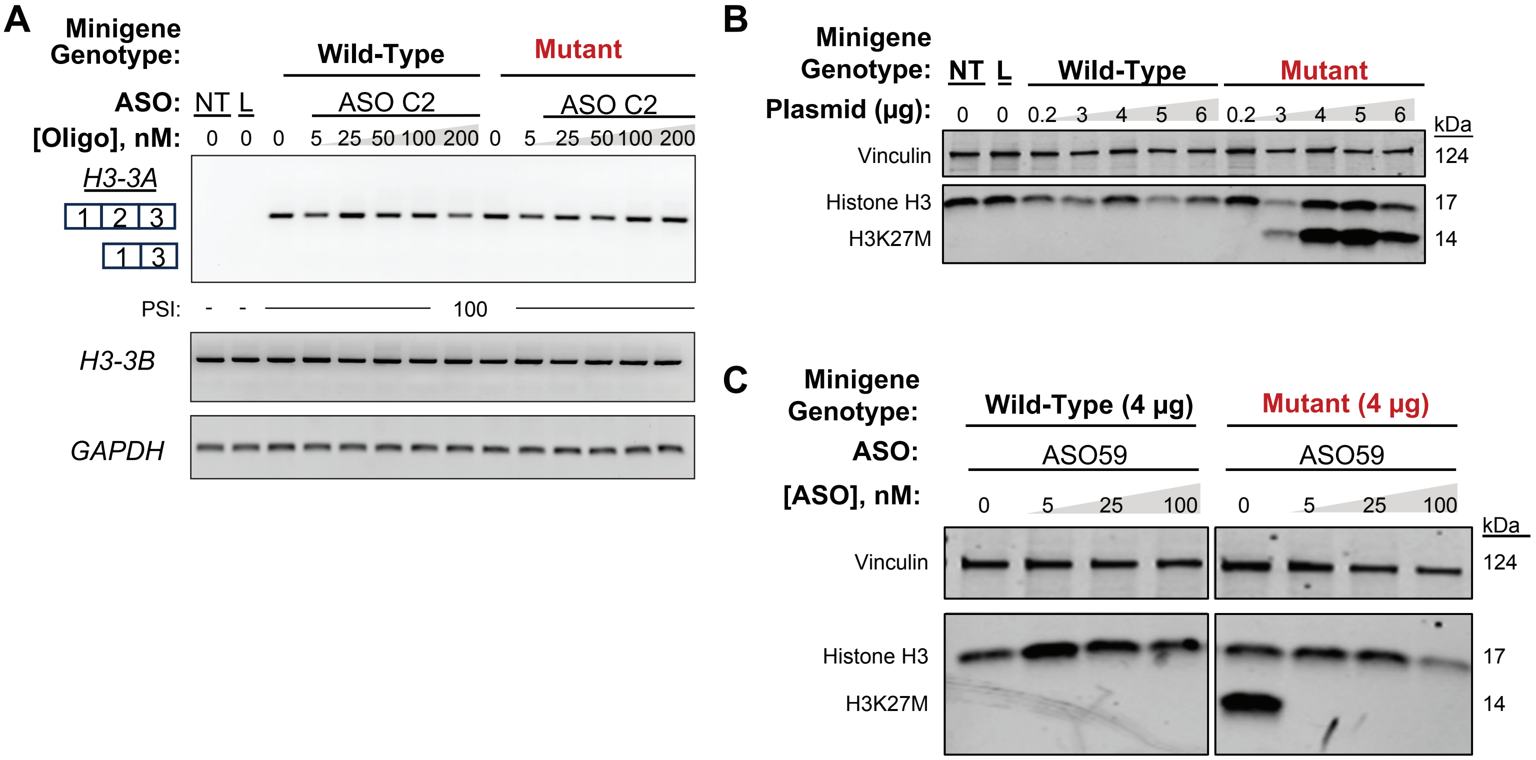
A
Minigene Genotype:
Wild-Type
Mutant
ASO: NT L
ASO C2
ASO C2
[Oligo], nM: 0 0 0 5 25 50 100 200 0 5 25 50 100 200
H3-3A
1 2 3
1 3
PSI: - - 100
H3-3B
GAPDH
B
Minigene Genotype: NT L Wild-Type Mutant
Plasmid (µg): 0 0 0.2 3 4 5 6 0.2 3 4 5 6
kDa
Vinculin 124
Histone H3 17
H3K27M 14
C
Minigene Genotype:
Wild-Type (4 µg)
Mutant (4 µg)
ASO:
ASO59
ASO59
[ASO], nM: 0 5 25 100 0 5 25 100
kDa
Vinculin 124
Histone H3 17
H3K27M 14

Figure S4

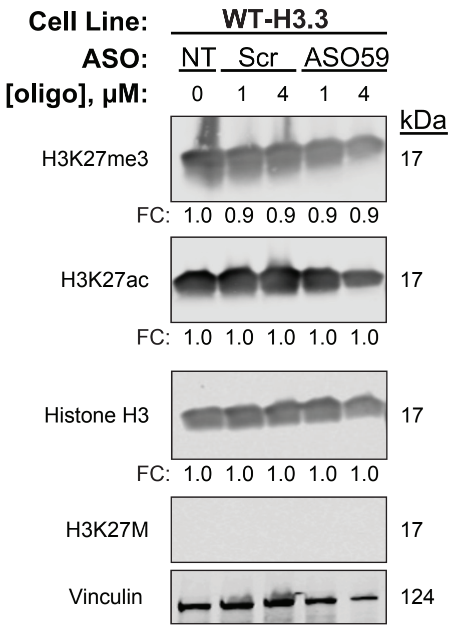
Cell Line:
WT-H3.3
ASO:
NT
Scr
ASO59
[oligo], μM:
0
1
4
1
4
kDa
H3K27me3
17
FC: 1.0 0.9 0.9 0.9 0.9
H3K27ac
17
FC: 1.0 1.0 1.0 1.0 1.0
Histone H3
17
FC: 1.0 1.0 1.0 1.0 1.0
H3K27M
17
Vinculin
124

Figure S5

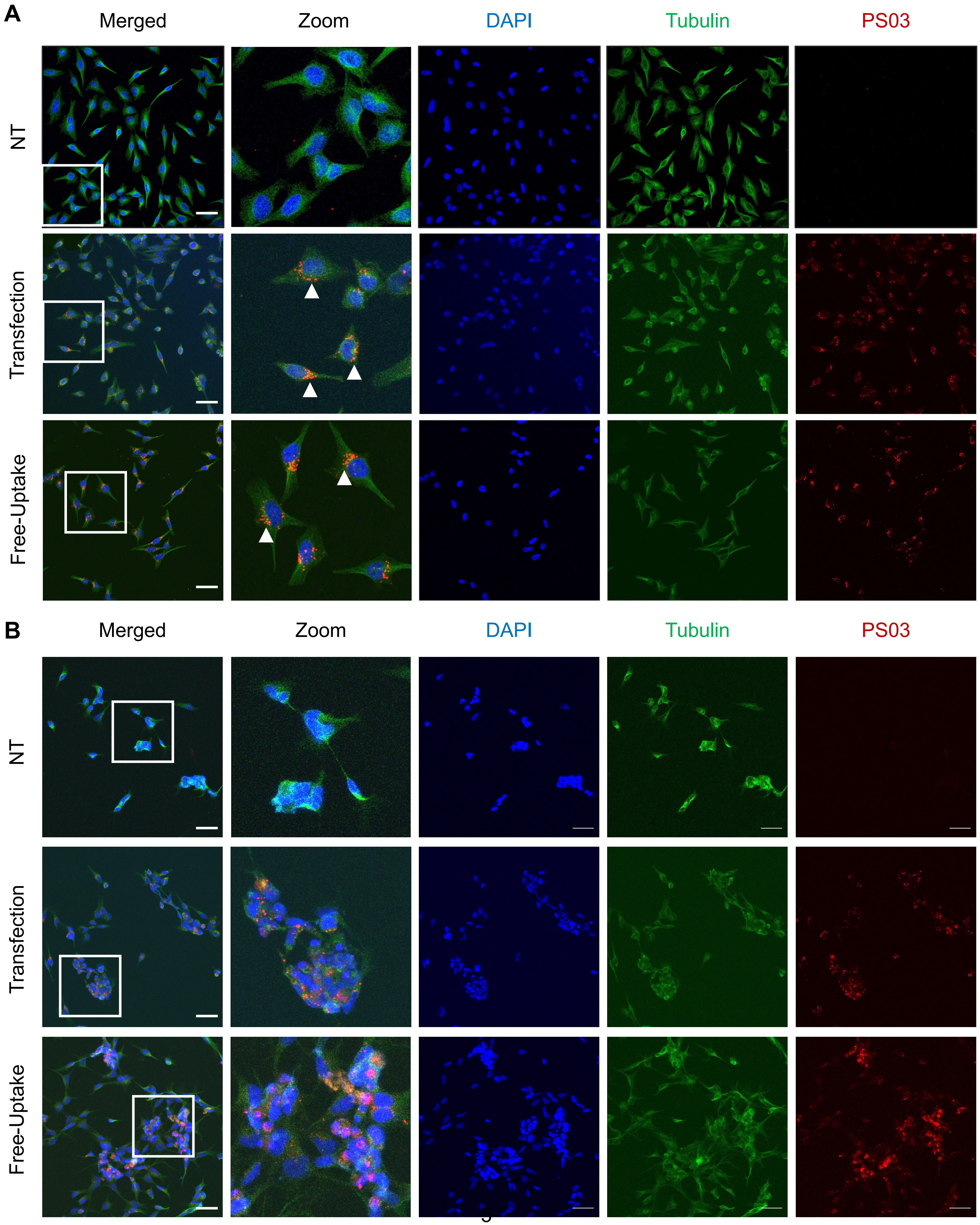
A
Merged
Zoom
DAPI
Tubulin
PS03
NT
Transfection
Free-Uptake
B
Merged
Zoom
DAPI
Tubulin
PS03
NT
Transfection
Free-Uptake

Figure S6

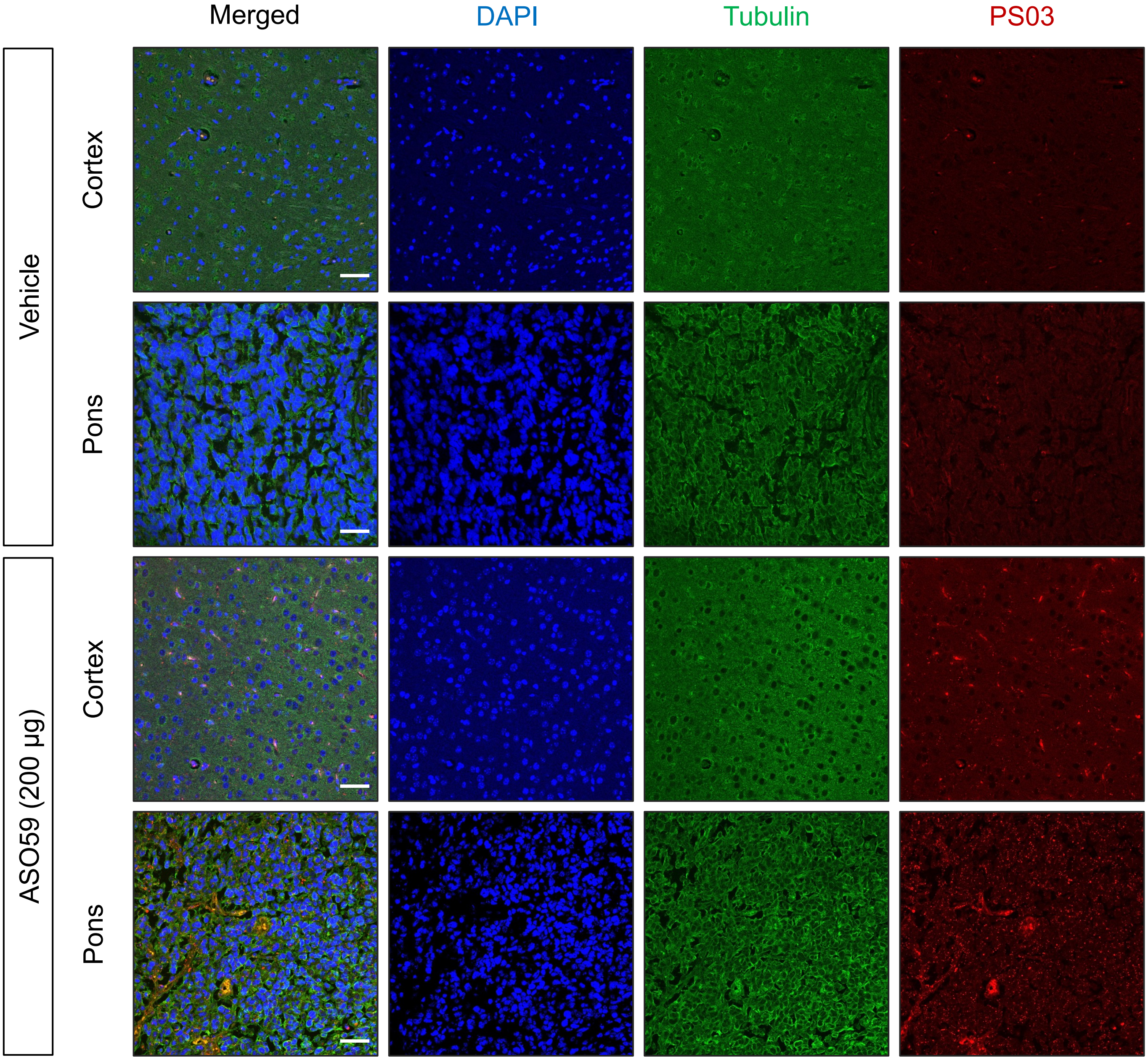
Merged
DAPI
Tubulin
PS03
Vehicle
Cortex
Pons
ASO59 (200 µg)
Cortex
Pons

Figure S7

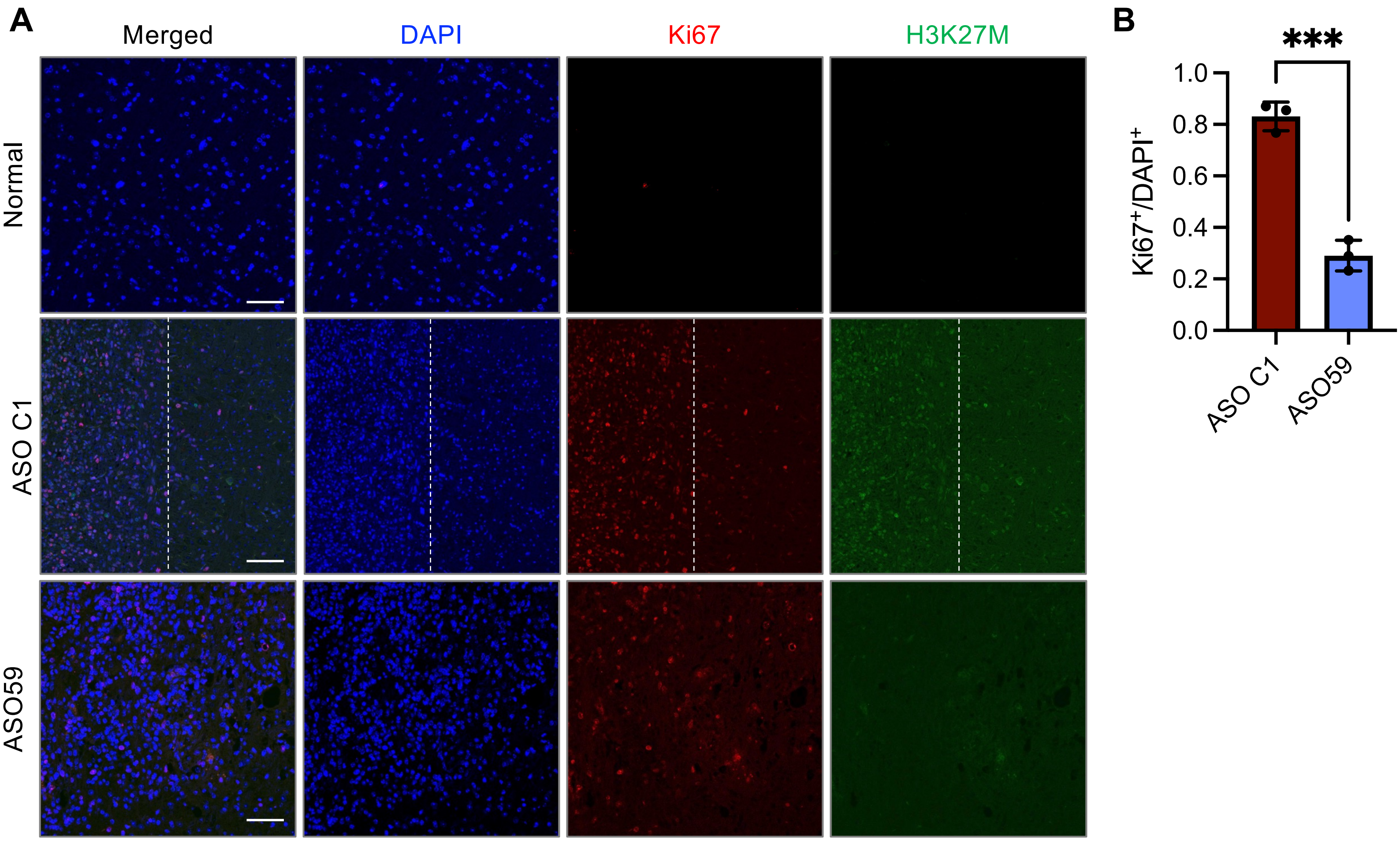
A
Merged
DAPI
Ki67
H3K27M
Normal
ASO C1
ASO59
B
***
1.0
0.8
0.6
0.4
0.2
0.0
Ki67+/DAPI+
ASO C1
ASO59

**Supplemental Table S1. List of ASOs.**

| ASO Name | Sequence (5’-to-3’) |
|---|---|
| ASO58 | TTTCCTGTTTTTTAATACCT |
| ASO59 | TTCCTGTTTTTTAATACCTG |
| ASO60 | TCCTGTTTTTTAATACCTGT |
| ASO61 | CCTGTTTTTTAATACCTGTA |
| ASO62 | CTGTTTTTTAATACCTGTAA |
| ASO63 | TGTTTTTTAATACCTGTAAC |
| ASO64 | GTTTTTTAATACCTGTAACG |
| ASO65 | TTTTTTAATACCTGTAACGA |
| ASO66 | TTTTTAATACCTGTAACGAT |
| ASO67 | TTTTAATACCTGTAACGATG |
| ASO68 | TTTAATACCTGTAACGATGA |
| ASO69 | TTAATACCTGTAACGATGAG |
| ASO70 | TAATACCTGTAACGATGAGG |
| ASO71 | AATACCTGTAACGATGAGGT |
| ASO72 | ATACCTGTAACGATGAGGTT |
| ASO73 | TACCTGTAACGATGAGGTTT |
| ASO74 | ACCTGTAACGATGAGGTTTC |
| ASO C1 | TTATTTCTGTATTCTATCCG |
| ASO C2 | TACAGAGACCTCCTTACTTA |

**Supplemental Table S2. List of Primers.**

| Primer Name | Sequence (5'-3') |
|---|---|
| T7 Forward | TAATACGACTCACTATAGGG |
| *H3-3A* Exon 1 Reverse | CCCCCTTCTCCTTCGGCTGG |
| *H3-3A* Exon 3 Reverse | AAGTCCTGAGCAATTTCTCG |
| *H3-3A* Exon 1 Forward (for splicing) | CCGAGCTCCAGCCGAAGGAG |
| *H3-3A* Exon 4 Reverse (for splicing) | AAGCACGTTCTCCACGTATGC |
| *H3-3B* Exon 1 Forward | GGTTTTTCGCTCGTCGACTG |
| *H3-3B* Exon 4 Reverse | GGTTGGTATCTTCGAACAGA |
| *GAPDH* Forward | AATCCCATCACCATCTTCCA |
| *GAPDH* Reverse | TGGACTCCACGACGTACTCA |

**Supplemental Table S3. *In Vivo* Data from H3.3K27M-Altered DMG PDX Samples.**
Please see separate Excel file.

**Supplemental Table S4. Summary Statistics of H3.3K27M-Altered PDX Survival.**

| | Total # of Mice | # Male Mice | # Female Mice | Median Survival (days) | Mean Survival (days) |
|---|---|---|---|---|---|
| **Vehicle** | 7 | 4 | 3 | 26 | 27 |
| **ASO C1** | 7 | 3 | 4 | 26 | 27 |
| **ASO59** | 7 | 3 | 4 | 43 | 43 |

**Supplemental Table S5. Survival Analysis of H3.3K27M-Altered PDX.**

| | **Log-Rank (Mantel Cox)** | | | **Gehan-Breslow-Wilcoxon** | | |
|---|---|---|---|---|---|---|
| | Chi-Square | P-value | Summary | Chi-Square | P-value | Summary |
| **Vehicle vs ASO C1** | 0.09157 | 0.7622 | ns | 0.01627 | 0.8985 | ns |
| **Vehicle vs ASO59** | 14.12 | 0.0002 | *** | 12.11 | 0.0005 | *** |
| **ASO C1 vs ASO 59** | 11.26 | 0.0008 | *** | 10.3 | 0.0013 | ** |
